# Pre-neuronal processing of haptic sensory cues via dispersive high-frequency vibrational modes

**DOI:** 10.1101/2022.06.15.496141

**Authors:** Yu Ding, Yurii Vlasov

## Abstract

Sense of touch is one of the major perception channels. Neural coding of object textures conveyed by rodents’ whiskers has been a model to study early stages of haptic information uptake. While high-precision spike timing has been observed during whisker sweeping across textured surfaces, the exact nature of whisker micromotions that spikes encode remains elusive. Here, we discovered that a single micro-collision of a whisker with surface features generates vibrational eigenmodes spanning frequencies up to 10KHz. While propagating along the whisker, these high-frequency modes can carry up to 80% of shockwave energy, exhibit 100X smaller damping ratio, and arrive at the follicle 10X faster than low frequency components. The mechano-transduction of these energy bursts into a time-sequenced population spike trains may generate temporally unique “bar code” with ultra-high information capacity. This hypothesis of pre-neuronal processing of haptic signals based on dispersive temporal separation of the vibrational modal frequencies can shed light on neural coding of haptic signals in many whisker-like sensory organs across the animal world as well as in texture perception in primate’s glabrous skin.

**Significance Statement:** Understanding how the outside world is encoded in neurons spikes in sensory organs and how these neural codes contribute to perception remains elusive. Using a model system - a whisker of a mouse - we discovered that tiny whisker vibrations induced at the whisker tip by collisions with external objects generate a time series of energy bursts. This creates a temporally unique “bar code” of a time-sequenced population spike trains with ultra-high information capacity. We hypothesize that such a “pre-neuronal processing” of touch events into time-coded spikes can provide a conceptual link to understand neural coding in many whisker-like sensory organs across the animal world as well as in texture perception in primate’s glabrous skin.

## INTRODUCTION

Understanding how the outside world is encoded in neurons spikes in the mammalian brain and how these neural codes contribute to perception remains elusive. To get insights in sensory perception we use here a model sensory system - a whisker of a mouse that is the main sensory input channel to this mostly nocturnal animals. Rodents use their mystacial vibrissae (whiskers) to explore the environment and gather information about nearby objects. Active back and forth rhythmic movement of whiskers over objects (whisking) is used to produce a neural representation of a complex tactile scenes enabling rodents to locate objects (*1–3*), to perform object recognition (*4*), to guide their locomotion along walls (*5, 6*), and to navigate in their natural habitat of dark burrows and tunnels (*7*). Whiskers themselves are not innervated. Instead, numerous collisions of moving whiskers with objects produce time-varying forces at the whisker follicle (*8, 9*) that are picked by mechanosensitive nerve endings (mechanoreceptors) of the primary sensory neurons (*10*) in the trigeminal ganglion (TG) (*11*) to produce a barrage of neural discharges that is then directed to thalamo-cortical circuits generating haptic perception.

While the primary sensory neurons can perform complex computations (*12–16*), it is well recognized, however, that transduction of whisker collisions into forces at the whisker base is by itself a nonlinear process that effectively contributes to coding of haptic features and is often dubbed as a “pre-neuronal processing” (*17, 18*). Biophysical properties of the whisker (*8, 9*) as its length (*4*), taper(*19*), intrinsic curvature (*20*) and central medulla (*21*), as well as the surrounding follicle tissue (*22*), all contribute to complex transduction over large range of frequencies up to thousands of Hertz.

Transduction of quasi-steady-state whisker bending moments and forces into spike rate coding for azimuthal and radial localization is relatively well understood (*3, 8, 15, 23, 24*). Perception of texture is a more complex process that is likely mediated by high frequency whisker micromotions (*18, 25, 26*). One of the proposed transduction hypothesis considers an individual whisker as a resonant oscillator whose fundamental frequency depends on the whisker length (*26, 27*). Considering all whiskers across a mystacial pad having different length, hence driving different neuronal responses, such differential frequency tuning can be the origin of texture discrimination. An alternative “stick-slip” hypothesis (*18, 28–31*) considers a time sequence of sticking and slipping events when whisker is rubbed over a rough surface. Such “stick-slip” patterns of whisker movements result in fast variation of angular velocity (*32*), and associated angular acceleration peaks (*28, 30*) that are shown to drive cortical activity (*29*). Neuronal spiking is found to be highly synchronized to stick-slip acceleration peaks with less than 2ms jitter in primary somatosensory cortex (*29, 31*), sub-milliseconds in thalamus (*33*), and as small as 10μs in primary afferents (*34*). These characteristic times correspond to frequencies of whisker micromotions extending beyond 10-500Hz that are typically studied with high-speed video recordings with subsequent image analysis (*35*). What aspects of such fast and complex spatiotemporal micromotions of a whisker are contributing to generation of these precisely timed spikes? Knowing “what” is encoded in neurons spikes may shed light on “how” this information is decoded and processed in thalamo-cortical neural circuits to produce tactile perception.

Here, using highly sensitive acoustical methods we performed a systematic study of whisker micromotions and identified a regular series of higher order vibrational modes (up to 30) spanning frequencies up to 10KHz. We formulate a dispersive vibrational transduction hypothesis that considers a whisker a dispersive pre-neuronal processor that transforms high-frequency micromotions into an efficient temporal code with ultra-high Kb/s bandwidth.

## RESULTS

### Whisker micromotions are segregated into distinct transverse vibrational modes extending to 10KHz

While it is possible to detect whisker shape distortions using fast video recordings at frequencies up to 300Hz (*35*) and even measure the propagation speed of a shockwave (*36*), studies of micromotions at frequencies beyond a few hundred Hz are challenging. Instead, here we utilized acoustical methods (Methods, Fig.S1) that provide calibrated (Fig.S2) measurements of the forces at the whisker base covering over 50dB dynamic range over 10KHz bandwidth. Figure 1A shows a typical voltage signal recorded within the first few milliseconds after a mouse C1 whisker slips off a metal pole. Analogous data recorded with a B1 rat whisker are shown in Fig.1D. The results of the spectral analysis of this complex oscillatory pattern using a continuous Complex Morlet Wavelet (CMW) transformation (Methods) are shown in Fig.1B (mouse) and Fig.1E (rat). Vibrations are segregated into a distinct regular pattern of vibrational bands that are highly reproducible (SI Movie.1). The spectrogram cross-sections taken at 1ms delay (Fig.1C and Fig.1F, black curves) show regular peaks with 30dB contrast ratio that extend up to 10KHz. With consecutive trimming of the whisker distal end the vibrational bands are shifting to higher frequencies and their spectral separation is increasing (Fig.2C and Fig.2F).

**Figure 1.**
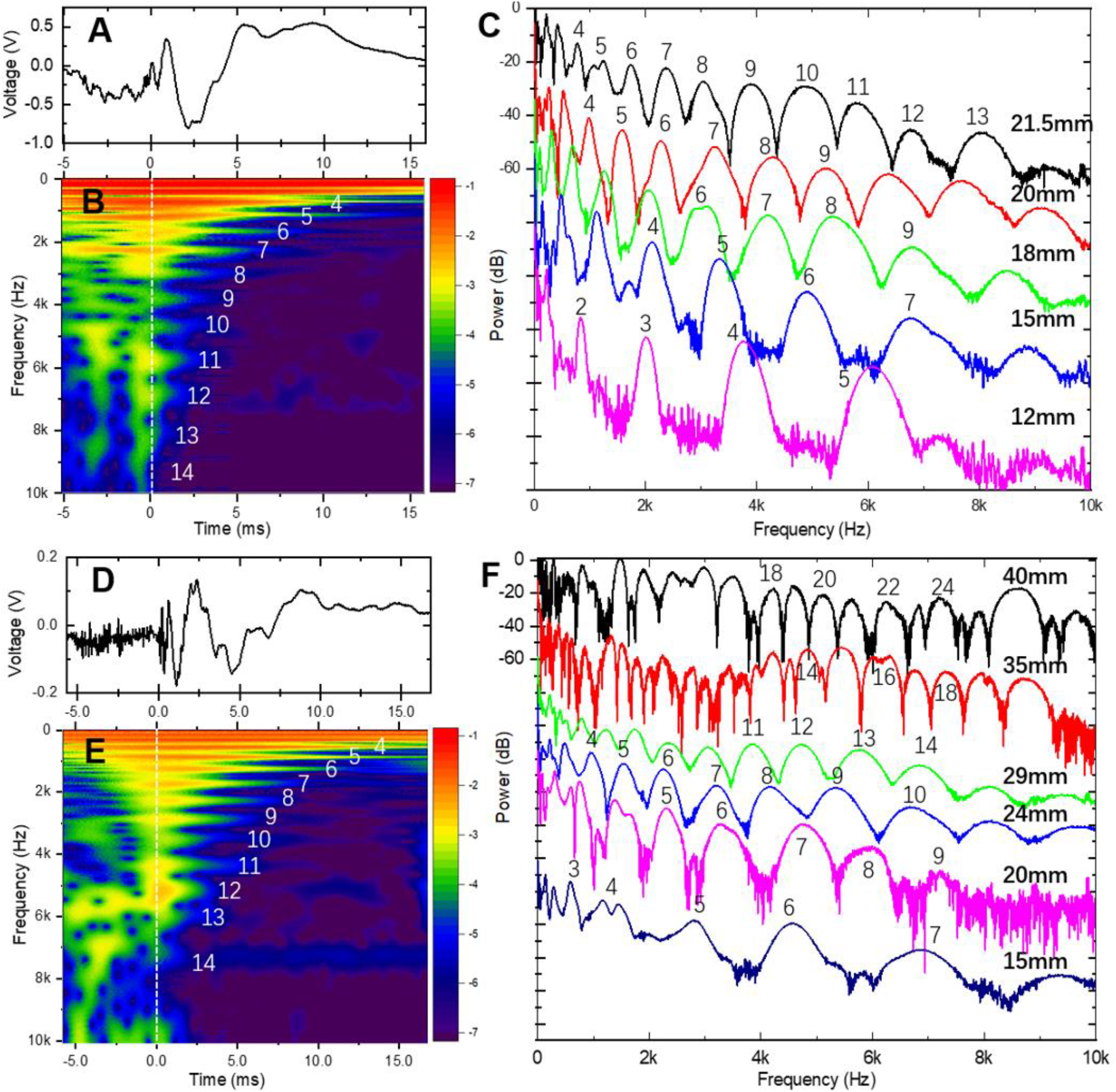
Whisker micromotions segregate into distinct vibrational bands up to 10KHz. **(A)** Voltage trace recorded when 21.5mm long C1 mouse whisker slips-off the pole in a concave forward direction. Sweep rate 3.5Hz with position of the pole 10mm away from the whisker distal end. (**B)** CMW spectrogram of the voltage trace in A). Color scheme represents the transform absolute magnitude in logarithmic scale. Vibrational bands are numbered starting from the fundamental mode. (**C)** Amplitude of the CMW transforms taken at 1ms time delay from the slip-off for the C1 mouse whisker that is consecutively trimmed. Curves are shifted vertically for clarity. (**D)** Same as (A) for the 29mm long B1 rat whisker. Sweep rate 3.5Hz with position of the pole 2mm away from the whisker distal end. (**E)** CMW spectrogram of the voltage trace in D). (**F)** Same as (C) for the B1 rat whisker that is consecutively trimmed.

**Figure 2.**
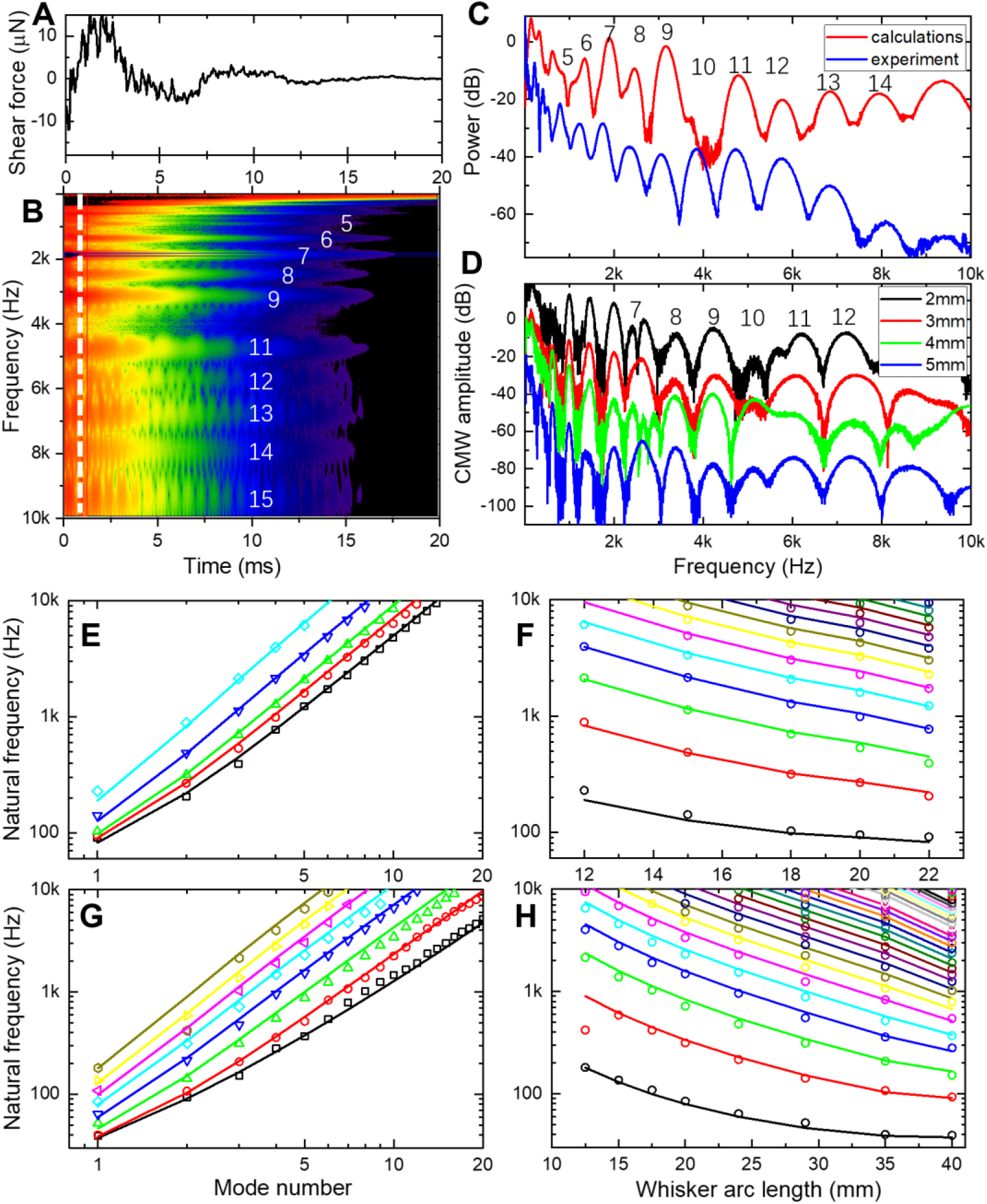
Vibrational bands are Euler-Bernoulli transverse eigenmodes. **(A)** Calculated shear force at the whisker base for a truncated whisker with *S*_*tot*_=29mm, *p*=1000kg/m^3^, *E*=2GPa, damping α = 430 rad/s. Whisker is touching the pole at 2mm from the tip. (**B)** CMW spectrogram of the voltage trace in A). (**C)** Normalized spectrum of the calculated CMW (red) taken at 1ms time delay from the slip-off in A), compared to normalized experimental power spectrum (blue) of B1 rat whisker trimmed to 29mm from Fig.1F. (**D)** Series of calculated CMW spectra for different position of the pole relative to the tip of the whisker (2, 3, 4, and 5mm from the tip). Whisker parameters are the same as in A). (**E)** Log-log plot of modes natural frequencies for the C1 mouse whisker that is consecutively trimmed. (**F)** Same results as in F), replotted in semi-log plot of modes frequencies vs whisker length. (**G)** Log-log plot of modes natural frequencies for the B1 rat whiskers that is consecutively trimmed. (**H)** Same results as in G) replotted in semi-log plot of modes frequencies vs whisker length.

These modes, numbered from the lowest frequency fundamental mode (n=1 at 63Hz for Fig1.B) up to the highest observed frequency band (n=14 at 9430Hz for Fig.1B), can be assigned to transverse vibrational eigenmodes of a freely vibrating truncated conical Euler-Bernoulli beam model without dissipation (*26, 27, 37*) described by the equation:

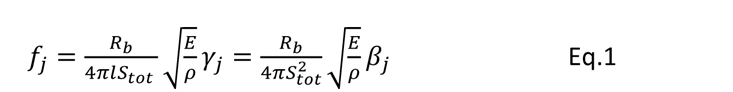

*f*_*j*_ where *f*_*j*_ is the linear natural frequency of the j-th mode, *R*_*b*_, *ρ*, *E* represent whisker radius at the base, its density, and Young’s modulus. The total arc length of the whisker before truncation is *S*_*tot*_, and the arc length after truncation at the distal end is *l*. The variables *β*_*j*_ (*γ*_*j*_) (SI Table.S2) are the dimensionless coefficients for full (truncated) whisker that depend on the eigenmode number as well as on boundary conditions (*38*).

### Vibrational bands are Euler-Bernoulli transverse vibrational eigenmodes

Using analytical dynamic model of whisker vibrations (*37*) (Methods) a time-dependent shear force at the base of the whisker after the slip-off is calculated and presented in Fig.2A. Corresponding CMW spectrogram in Fig.2B shows a series of individual vibration bands with frequencies closely matching experimentally observed values. Not only the frequency of the band peaks (Fig.2C), but also the band shapes are well predicted by theoretical calculations. Since modes phases and amplitudes depend strongly (Fig.2D) on specific location of a pole contact along the whisker length (*37*) that can explain a noticeable difference for bands 5 and 10. Figure 2E shows the evolution of extracted natural undamped (Methods, Fig.S5) frequencies of eigenmodes for the C1 mouse whisker consecutively trimmed from the initial 21.5mm arc length down to 12mm. Figure 2G shows analogous results for the B1 rat whisker. All 49 experimental data points in Fig.2E were fitted simultaneously using the Eq.1 model of with just a Young’s modulus as a single adjustable parameter. The best fit yields *E* =3.9583 GPa (Levenberg-Marquardt, standard error (SE) 0.0029, Pearson correlation coefficient *r*_*p*_=0.9989, p-value P_p_<0.0001). Similar data fitting for 107 data points for the B1 rat whisker (Fig.2G) yields *E* =1.7340 GPa (SE 0.0042, *r*_*p*_= 0.9890, P_p_<0.0001). Both of these numbers are within the previously reported ranges for rats (*21*) and mice whiskers (*37*), extracted, however, with higher accuracy. Same data are replotted in Fig.2F and Fig.2H on a semi-log plots to illustrate dependence of each mode natural frequency on the whisker length. Good fit for all whisker lengths indicate that contributions of Young’s modulus variation along the whisker length (*39*), density variation due to central medulla (*21*), or deviation of a whisker taper from the ideal conical shape (*40*) are relatively small.

### Modal damping is frequency dependent

Once the whisker is slipping off the pole and is freely vibrating in the air, the vibrational modes are decaying in time due to various energy dissipation channels including dissipation within the whisker, damping in the follicle, and friction in the air. Classical mechanics of the damped harmonic oscillator considers a damping ratio *ε* = 1⁄*ω*_*n*_*τ*, where *ω*_*n*_ is a natural frequency of the vibration mode and *τ* is temporal energy dissipation rate. Damping ratio quantifies how many oscillation cycles the vibration mode is going through prior to being dissipated. Historically, (*26, 27, 36, 37*) a linear damping model with a frequency-independent damping ratio *ζ* is typically assumed (*35, 37*), which results in identical temporal decay of each of vibrational bands in Fig.2B. This assumption, while convenient, is, however, unphysical as it violates causality.

In a striking contrast to this model, the experimental spectrogram of Fig.1B (and Fig.1E) exhibits increasingly fast time decay *τ* of the wavelet amplitude for higher frequencies indicating strong frequency dispersion of damping. A set of temporal cross-sections of CWM spectrograms taken at the eigenmodes central frequencies shows (Fig.3A, mouse and Fig.3C, rat) exponential temporal decay spanning 30-40dB in amplitude with a slope rapidly increasing with the mode number. The damping ratio *ζ* extracted (Methods) from the decay time constant *τ* of the fundamental mode wavelet temporal cross-section for the untrimmed mouse whisker (Fig.3B, black circles) is relatively large (*ζ*=0.3) indicating that fundamental mode is barely oscillating – condition called “heavily damped”. In contrast, while the high frequency modes exhibit much faster decay times *τ*, the resulting damping ratio *ζ* is, in fact, decreasing (Fig.3D) since higher order modes have higher natural frequency *ω*_*n*_. The damping ratio of modes with *ω*_*n*_ above 1KHz is reduced below 10^-2^ signifying that higher order modes are going through many hundreds of oscillation cycles before they lose their energy.

**Figure 3.**
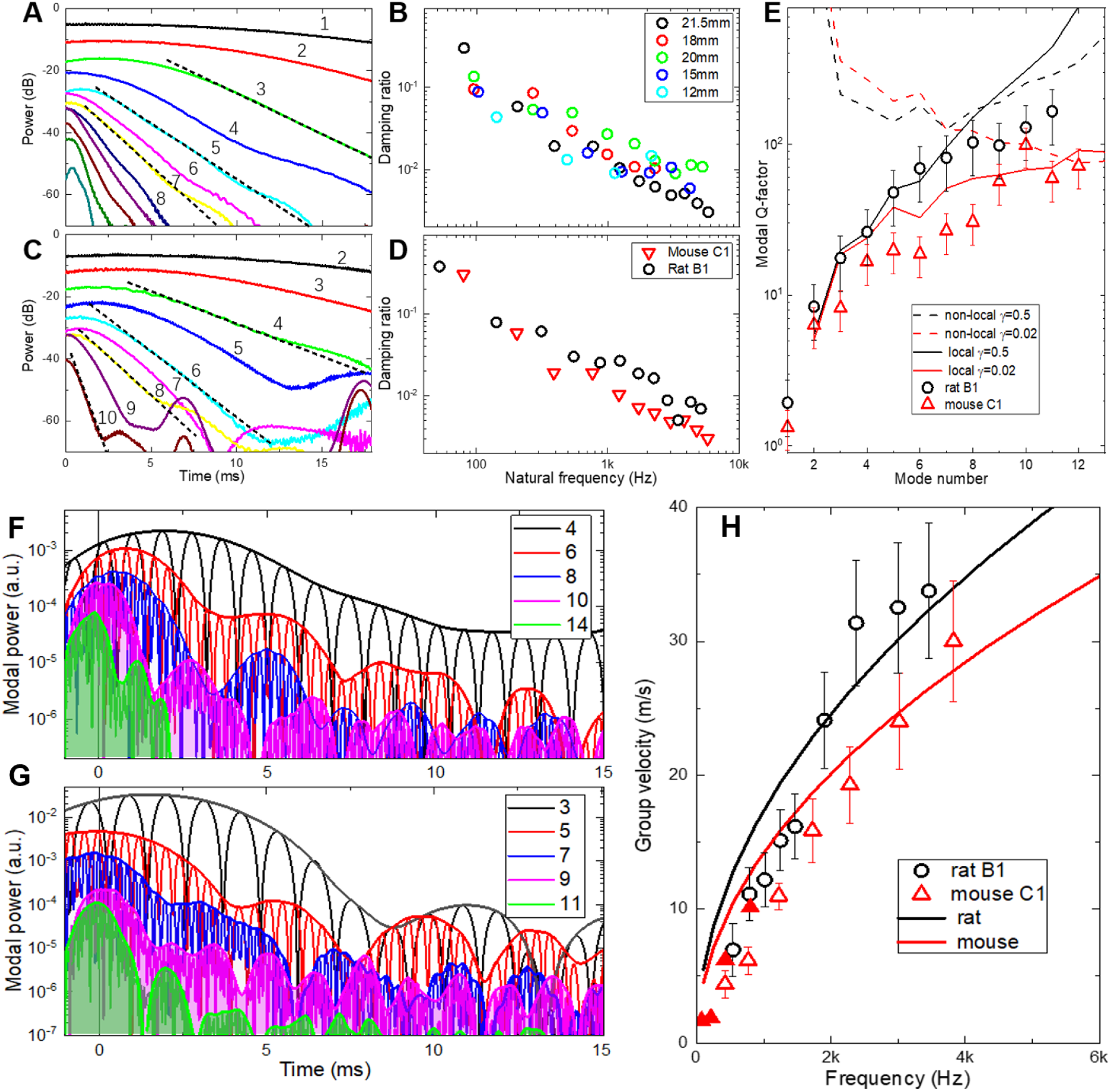
Damping and group velocity of vibrational eigenmodes are frequency dependent. **(A)** Set of cross-sections of the CMW spectrogram of Fig.1B taken at the central frequencies of corresponding vibrational modes. Dotted lines are linear fits to extract decay time and damping ratios. **(B).** Damping ratios for a consecutively trimmed C1 mouse whisker. **(C)** Same as A) for the consecutively trimmed B1 rat whisker. **D)** Comparison of damping ratios for the mouse (black circles) and rat (red triangles) whiskers. **(E)** Comparison of experimental modal Q-factors with non-locally reacting (dashed lines) and local (solid lines) models with viscous (γ=0.05, black) and non-viscous (γ=0.5, red) damping. **(F)** Set of FFT bandpass filtered power traces for different modes for the rat B1 whisker trimmed to 29mm arc length. **(G)** Same for the mouse C1 whisker with trimmed 20mm arc length. **(H)** Modal group velocity for the C1 mouse (red triangles) and B1 rat (black circles) whiskers. Filled triangles correspond to delays extracted from the envelope maxima. Solid lines are dispersion relations for the Euler-Bernoulli rod model of Eq.3 calculated for mouse (red) and rat (black) whisker parameters.

The dependence is roughly linear in a log-log scale indicating the inverse dependence of the damping ratio with frequency. Moreover, similar trend is observed for the same whisker that is consecutively trimmed to increasingly shorter arc lengths (Fig.3B) indicating that possible contributions of non-uniformity in whisker mechanical properties (tapering, Young’s modulus, density, etc.) to damping is relatively small. Analogous inverse frequency dependence is observed also for the rat B1 whisker trimmed to 29mm arc length (Fig.3C, D) indicating similar damping mechanisms.

For lightly damped oscillator, complex eigenfrequency of the vibrational mode can be expressed in terms of the undamped natural frequency *ω*_*n*_ and the modal Q-factor that is 1⁄2*ε* and is proportional to the ratio of the total energy stored divided by the energy lost per cycle. The Q-factors are presented in Fig.3E for the C1 mouse (black circles) and the B1 rat (red triangles) whiskers. To identify various types of damping we used a simple method that has been proposed for identification of various types of damping matrix (*41*) that is based on evaluation of a parameter *γ* that is a fraction of a characteristic relaxation time of the modal damping function compared with the natural periods of the vibration. Solid and dashed lines in Fig.3E represent theoretical modal Q-factors for different *γ* values taken from Ref.(*41*) for an exponential relaxation function. Comparison with the non-locally reacting damping models (dashed lines in Fig.3E) and with locally reacting (solid lines) the experimental data support local damping models. While for the mouse whisker *γ* is as small as 0.02, the damping is essentially viscous. Since for the rat whisker *γ* is closer to 0.5 its damping matrix has potential contribution of non-viscous asymmetric terms.

### Modal group velocity is frequency dependent

When the shockwave is generated by a rapid application of a force to a whisker during interaction with the object, it does not reach the whisker base immediately but propagates with a finite speed. Since the shockwave is an amplitude modulated wave packet, it propagation is described by a group velocity *ν*_*g*_ rather than phase velocity. The shockwave group velocity has been measured (*36*) and calculated (*37*) to be of the order of 5-10m/s that provides a few ms delay for the vibrational energy to reach the follicle (*42*). However, as it follows from previous sections, the shockwave is a superposition of multiple eigenmodes with their energy distribution dependent on where (Fig.2D) and how fast the force is applied (*37*). Since eigenmodes in a truncated conical beam interacting with a pole remain mutually orthogonal (*37*) even in the presence of viscous or non-viscous damping (*41*), the power of each eigenmode *U*_*j*_(*t*) is evolving independently over time:

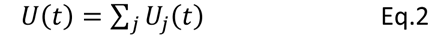

Therefore, not only the frequencies and damping ratios of the eigenmodes are different, but also their group velocities as well, since, in general, the transverse waves (as opposed to longitudinal or torsional waves) are dispersive (*43*). The group velocity *ν*_*g*_ of the transverse wave propagating in an infinite Euler-Bernoulli rod with a constant radius *R* is:

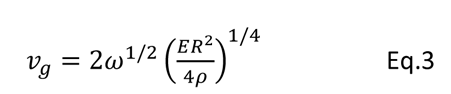

Spectral analysis based on FFT bandpass-filtering (Methods, SI Fig.S5) is shown for several higher order eigenmodes for the B1 rat (Fig.3F) and the C1 mouse whisker (Fig.3G). The maxima of the power envelopes of the lower order modes in Fig.3F, G are delayed by a few ms with respect to maxima of the higher order modes indicating normal group velocity dispersion of Eq.3 (see also Fig.3A,C, and Fig.S5B). Group velocity calculated from the arrival time of the envelop maxima for a C1 whisker is plotted in Fig.3H (solid red triangles) showing group velocity of 1.7m/s for the fundamental mode (74Hz) increasing to 10m/s for the 4^th^ mode (800Hz) (see also Fig.S5D).

Besides the apparent delay of the envelope maxima, a periodic modulation of the envelope amplitude is observed in Fig.3F, G with the beating period decreasing with the increase of the mode number from over 5ms for lower order modes to less than 1ms for higher frequency modes. We hypothesize that this periodic modulation is the result of the modes wave packets bouncing back and force due to strong reflections at the whisker base (reflection coefficient 0.61) and at the tip (over 0.99) owing to strong impedance mismatch between the keratin whisker (1.7×10^6^ Rayl), metallic microphone at the base (14×10^6^ Rayl) and air at the tip (4×10^2^ Rayl). Fig.3H represents the group velocity dispersion for both rat (black circles) and mouse (red open triangles) whiskers. Solid lines in Fig.3H show group velocity calculated using Eq.3 model using parameters from the Fig.2 fits. Even though the model assumes a non-tapered cylindrical whisker, the overall trend and the magnitude of the group velocities are well reproduced.

Therefore, not only higher order modes have 100X lower damping ratios, but they also propagate at much faster group velocity of 20-30m/s. This implies that higher order modes propagate fast enough to the whisker base that they still might have enough energy to stimulate mechanoreceptors at the follicle, while lower order modes arrive later with much of the energy dissipated during travel.

### Higher order modes carry up to 80% of the shockwave vibrational energy

Interaction of a whisker with an isolated pole considered above is a relatively rare situation for rodents navigating in natural environment. To study much richer repertoire of whisker micromotions related to the natural stimuli, additional experiments were performed using the same experimental setup (Fig.S1), but with a pole replaced with a sandpaper of different grit numbers (Fig.S4 and SI Movie.2). Figure 4 represents typical voltage traces and CMW spectrograms for the same B1 rat (Fig.4A,B) and the C1 mouse (Fig.4D,E) whiskers. Transient short-lived vertical streaks in the CWM spectrograms are observed, representing fast events during whisker swiping across the textured surface that generate energetic shockwaves with prominent high frequency components. The frequency spectrum of each streak shows a series of vibrational bands with modal frequencies consistent with the whisker arc length shorter by 2-3mm corresponding to the portion of the whisker tip attached to the surface during sweeping.

**Figure 4.**
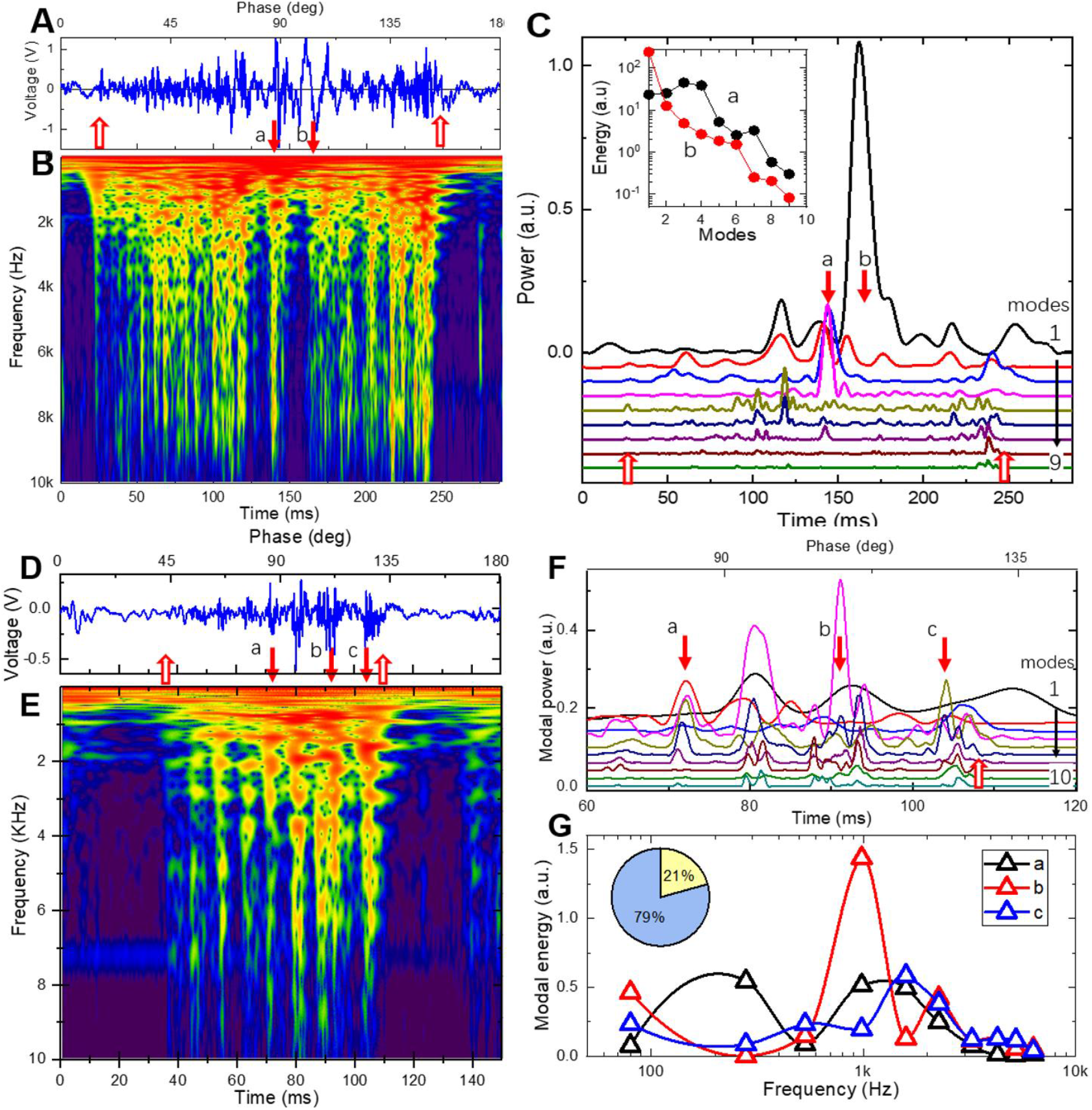
Texture induced high-frequency eigenmodes carry up to 80% of shockwave energy. **(A)** Voltage trace for a single cycle of a CF sweep of the 40mm long B1 rat whisker over a P60 grits sandpaper located 35mm from the base. Upward open arrows indicate the first touch (26ms) and the last detachment (250ms) events. **(B)** Complex Morlet wavelet transformation spectrogram of the voltage trace in A). Color scheme represents the transform absolute magnitude in a logarithmic scale. Events a and b shown by the downward arrows are discussed in the text (see also SI Movie.4). **(C)** A set of FFT bandpass filtered power traces for the 1 through 9 modes for the voltage trace in A). Downward arrows indicate same events a and b as in B). Inset: Energy distribution of different modes for events *a* and *b*. **(D)** Voltage trace for a single cycle of a CF sweep of the 21.5mm long C1 rat whisker over a p60 grits sandpaper located at 18mm from the base. Upward open arrows indicate the first touch (36ms) and the last detachment (109ms) events. **(E)** Complex Morlet wavelet transformation spectrogram of the voltage trace in D). Events *a, b,* and *c* shown by the downward arrows are discussed in the text. **(F)** A set of FFT bandpass filtered power traces for the 1 through 10 modes for the voltage trace in D). Downward arrows indicate same events *a, b,* and *c* as in B). **(G)** Energy distribution of different modes for events *a, b,* and *c* as in B). Inset: Ratio of combined energy carried by modes below 1000Hz (yellow, 21%) and high frequency modes above 1000Hz (blue, 79%) for the event *b*.

Naturally, such streaks can be associated with shockwaves induced by “stick-slip” whisker motions believed to be processed by primary afferents and upstream cortico-thalamic circuits to produce texture sensation (*28–34*). While each streak is accompanied by fast whisker tip acceleration (discussed in detail below), however each streak event is very unique. Spectral power analysis in Fig.4C shows that the power distribution between eigenmodes differs substantially even for neighboring vibrational events. To illustrate this observation, we picked an event at 160ms (Fig.4A, event *b*) when mostly the fundamental mode is excited. In contrast, the event at 145ms (Fig.4A, event *a*) the power of the higher order modes 3 and 4 are much higher than that of the fundamental mode. Modal energy (Fig.4C, inset) calculated for 5ms integration window (Methods) shows fast exponential decay for event *b* resembling pole-induced free vibration case (Fig.S3E). In contrast, during the event *a*, the modes 3 and 4 are over 10X more energetic than the fundamental. Similar spectral power analysis for the mouse whisker (Fig.4F) reveals even more striking observation that not only the power of the fundamental mode is relatively small (events *b, c*) or even virtually negligible (event *a*), but also the power is quickly redistributed towards higher order modes. The modal energy analysis (Fig.4G) shows that for most of the events the fundamental and the first two higher order modes combined carry together only around 20% of the total vibrational energy (Fig.4G, inset) while modes at frequencies beyond 1000Hz take up to 80% of the shockwave energy. Therefore, each shockwave event is characterized by a unique distribution of energies between different modes as expected from Eq.2. Comparison of CWM spectrograms for several nominally identical CF swipes (SI Movie.4) illustrate that, while on average the distribution between swipes is similar, however unique events (e.g. event *a* in Fig.4A) are characteristic of a specific trajectory of a whisker tip over specific bumps and troughs of the textured surface. Moreover, in striking difference to interaction with the pole, for some of the events the vast majority of the shockwave energy is redirected to higher order modes at frequencies beyond 500Hz, well above frequencies typically studied with fast videography.

### High order modes are generated during high-speed collision

Although whisker interactions with a textured surface are much closer to interactions in the natural environment than swiping off the pole, however their complexity hinders detailed analysis. To get deeper insight, a set of experiments was performed with a whisker swiping against an aluminum grating with periodically spaced grooves (Methods). Fig.5A shows whisker shapes extracted from a post-processed (Methods) high-speed movie recorded at 1460fps during a 109ms-long single swipe of a mouse C2 whisker with 24.1mm arc length over three neighboring teeth of the grating (SI Movie.4). Interaction with each of a grating tooth (events 1, 2, and 3 in Fig.5A) can be divided into three consecutive phases: ”pinned”, “unpinned” and “collision”. First, the “pinned” or “stick” phase, is characterized by a whisker tip movement slower than the angular scan velocity (note crowding of whisker traces in the beginning of each event in Fig.5A). Once “stuck”, the whisker curvature *κ*_*p*_ gradually increases (Fig.5B) until the second “unpinned” or “slip” phase follows when the whisker tip is released from the tooth edge with large acceleration (up to 2×10^6^ deg/s in Fig.5C) and a sudden drop in curvature (Fig.5B). These pinned and unpinned phases are similar to a slip-off event observed during interaction with a pole in Figs.1-3 as both excite mostly lower order vibrations. The third “collision” phase is unique for the interaction with a grating (and with textured surfaces of Fig.4) as it represents the moment when the whisker tip at high velocity is colliding with the edge of a next grating tooth (next obstacle on a textured surface). This is accompanied by a sudden termination of the curvature drop (Fig.5B), abrupt deceleration (Fig.5C), and appearance of large amplitude transients in the vibrational power trace (Fig.5D) as well as excitation of high frequency eigenmodes (Fig.5E).

**Figure 5.**
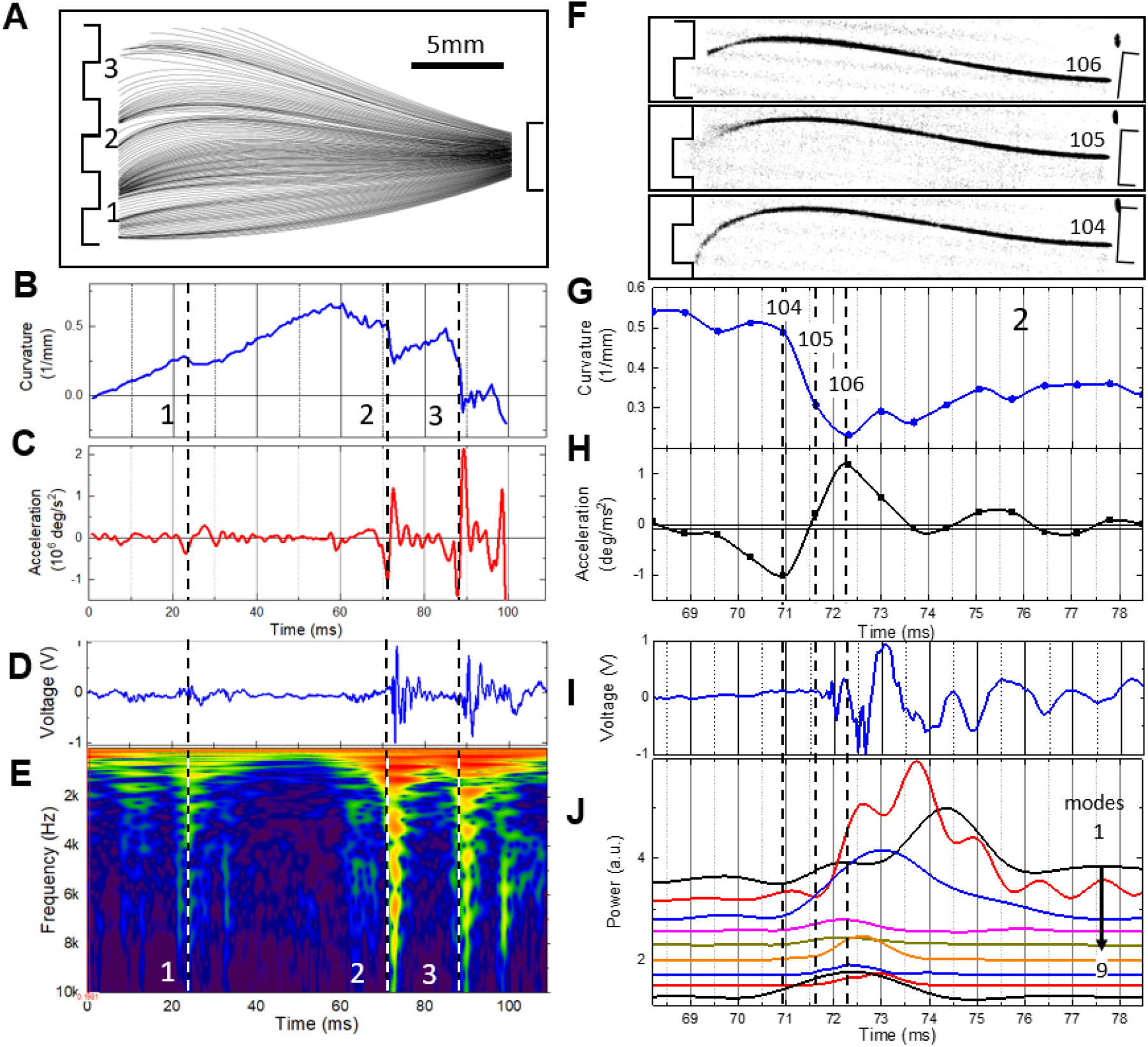
High-speed collisions induce dispersive shockwaves. **(A)** Shapes of a mouse C2 whisker with 24.1mm arc length extracted from consecutive 160 frames recorded at 1460fps during a single CB sweep over a grating. Whisker is traced for most of its arc length between 2mm from the base to 3mm from the tip. Events marked 1, 2, and 3 correspond to interaction with individual grating teeth. Note traces crowding at the beginning of each deceleration event (pinned phase), followed by acceleration (unpinned phase), and then deceleration when the whisker collides with the next tooth (collision phase). **(B)** Whisker curvature showing gradual increase during the pinned phase and distinct fast drops at the unpinned phase. **(C)** Angular acceleration of the whisker tip showing strong deceleration at the end of the pinned phase followed by large acceleration during the unpinned phase. **(D)** Voltage trace of a time-synchronized microphone recording. **(E)** CWM spectrogram of the voltage trace in D). **(F)** Three consecutive frames from the SI Movie.4 corresponding to pinned phase (frame #104), unpinned phase (frame #105), and collision phase (frame #106) of event 2. Note blurring at the whisker tip in the 105-frame corresponding to vibrations faster than 2KHz. **(G)** Whisker curvature for event 2. Time-synchronized frames are marked by symbols. Time position of frames in F) are shown by vertical dashed lines. **(H)** Angular acceleration of the whisker tip. **(I)** Time-synchronized recording of microphone voltage. **(J)** Envelop traces of the FFT bandpass filtered voltage for the first 9 eigenmodes. Note fast response of higher order modes and late arrival of lower order modes. Curves are shifted vertically by 0.25 for clarity.

Image of the frame #104 in Fig.5F corresponds to the end of the pinned phase characterized by a maximum curvature (Fig.5G) and largest deceleration (Fig.5H) indicating increase of a force applied to a whisker tip (hence increase of the stored elastic strain potential energy). This is followed by the frame #105 in Fig.5F that shows sudden whisker release (unpinned phase) when the force becomes large enough to overcome the static friction. This is accompanied by a sudden drop in whisker curvature (Fig.5G) with an offset indicative of an amount of kinetic energy released. Fast whisker vibrations corresponding to frequencies beyond 2000Hz show up as blurring at the whisker tip. Shockwave arrival time of eigenmodes of different frequencies at the whisker base

(Fig.5E) is consistent with the arc length from the base to the contact of 21mm (24.1mm total arc length minus 3mm protruding inside the groove). Therefore, even in the unpinned phase, the whisker tip is not vibrating freely, but is scratching the bottom of the grating groove. The very next frame #106 corresponds to a whisker tip collision with the edge of the next tooth, the end of a sudden curvature drop (Fig.5G), and the start of deceleration (Fig.5H). That corresponds to a “collision” phase when the kinetic energy of a moving whisker is converted into an energetic shockwave that starts to propagate along the whisker towards its base.

Time synchronized voltage trace (Fig.5I) shows that first signals appear at the whisker base at 71.7ms time (in-between frames #105 and #106). Spectral mode analysis of the voltage trace in Fig.5J reveals that, as in the case of interaction with a texture in Fig.4, the shockwave energy is efficiently converted to higher order modes. Note, that as in the case of interaction with the pole in Fig.S5C, the power of a second order mode (34% of total energy at 200Hz) exceeds the power of the fundamental mode (20% of total energy at 87Hz) consistent with prior observations (*42*). However, as opposed to interaction with the pole and consistent with observations of Fig.4 for a textured surface, the higher order modes carry substantial portion of a shockwave energy (i.e., 10% of total energy in the mode #9 at 3800Hz). These higher order modes arrive at the whisker base first, while the lower order modes are significantly delayed (e.g. fundamental mode #1 is delayed by 3ms) consistent with the group velocity dispersion of Fig.3H.

## DISCUSSION

### Whisker is a dispersive pre-neuronal processor

Results discussed above demonstrate that a whisker encounters a series of a 3-phase pinned-unpinned-collision events while interacting with the object (Fig.6A). The elastic energy stored in the whisker bending during the pinned phase is transformed into kinetic energy of a whisker movement during the unpinned phase. Whisker acceleration during pinned-unpinned phases is not enough to generate energetic high frequency modes above 500Hz. It is the whisker’s high-speed collision with an obstacle that converts the kinetic energy accumulated during pinned-unpinned phases into a vibrational shockwave. The shockwave is segregated into wave packets of individual vibrational eigenmodes (Fig.1 and Fig.2). The larger the kinetic energy (faster velocity) and the shorter the collision time, the broader is the frequency power spectrum of a shockwave and the more efficient is the excitation of higher order modes (*37*). Their excitation is, therefore, inefficient during slipping off the pole (less than 3% of shockwave energy, FigS3), but starts to show up during interaction with a grating (up to 30% of shockwave energy, Fig.5), and they can dominate in some of the events during interaction with a textured surface (up to 80% of shockwave energy, Fig.4). Being mostly orthogonal (Eq.2) even in the presence of damping, these modes are not exchanging energies while propagating. Instead, due to a modal dispersion of damping (Fig.3E) and of group velocity (Fig.3H), the shockwave, therefore, while propagating along the whisker, disperses with individual wave packets arriving at the base at different times and with different attenuation. Higher frequency modes above 500Hz (red oscillatory packet in Fig.6A) being fast and less damped arrive first, while slow and heavily damped low frequency modes (blue oscillatory packet in Fig.6A) are significantly delayed. A single collision event, therefore, is transformed into a time series of individual events at the whisker base as the eigenmode wave packets arrive.

**Figure 6.**
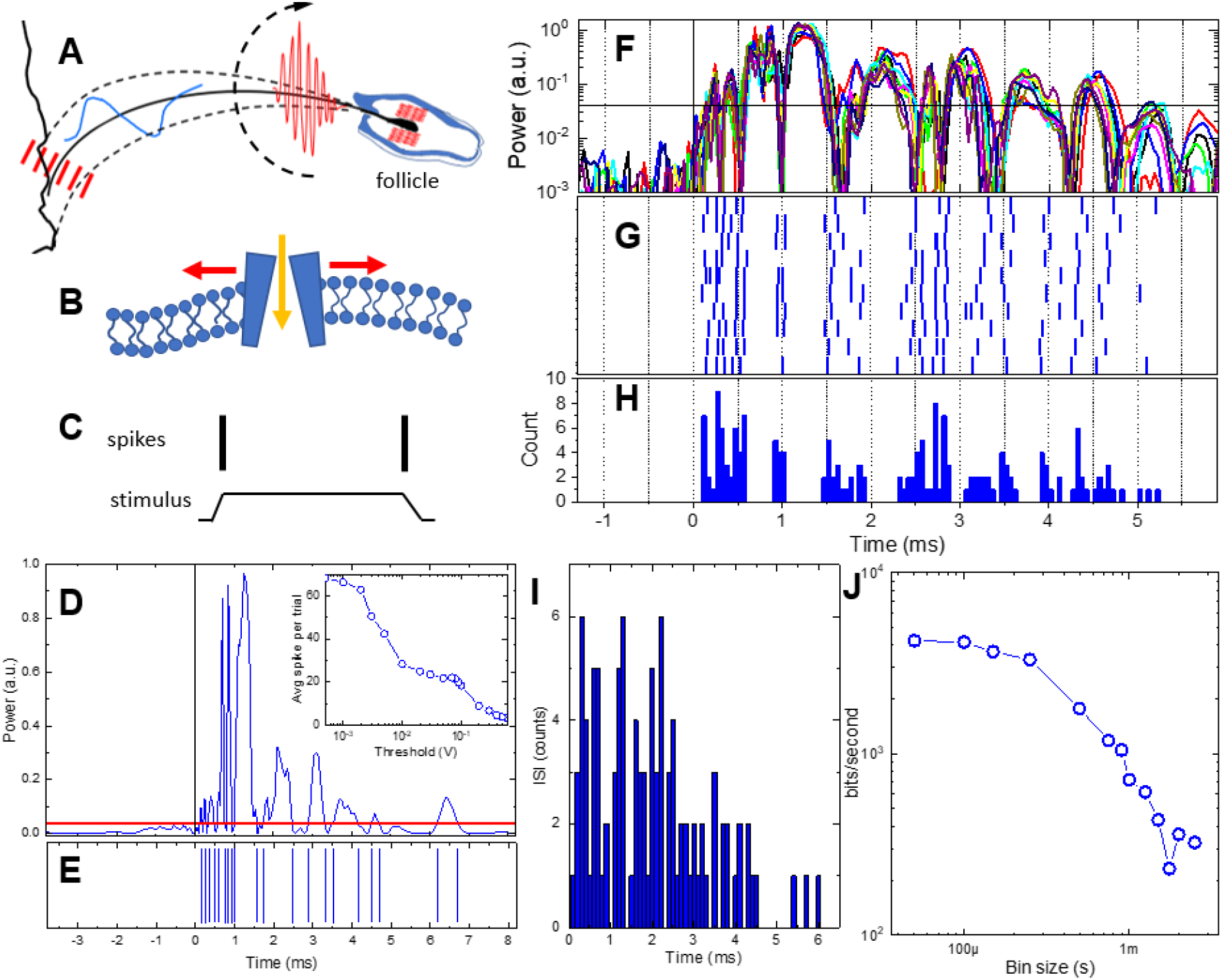
Dispersive Vibrational Transduction model predicts ultrahigh information capacity. **(A)** Schematic of a whisker with is a base inside a follicle. Red bundles in the follicle denote Rapidly Adapting (RA) mechanoreceptors. Whisker encounters a series of pinned-unpinned-collision events while being swept across the rough object. The shockwave generated during the collision phase (red bars on the left) propagates along the whisker towards the follicle. While propagating, the shockwave is dispersed, which results in higher frequency modes (red oscillatory packet) being fast and less damped arriving first, while slow and heavily damped low frequency modes (blue oscillatory packet) are significantly delayed. **(B)** Energetic wave packets arriving at the follicle induce opening of ion channels in piezo-sensitive membrane of a mechanoreceptor. **(C)** RA mechanoreceptors respond to fast stimulus by generating spikes (vertical black bars) at the very beginning and at the end of mechanical stimulation. **(D)** Power trace of event 2 (squared offset-corrected voltage of the same recording as in Fig.5J). The red line is a threshold at 0.04 level. **(E)** Artificial spike sequence generated by thresholding of D). Inset: Number of spikes per trial for different threshold levels. **(F)** Power traces (semi-log scale) of 10 nominally identical whisker sweeps corresponding to event 2 of Fig.5 aligned to the collision time (see also SI Movie 5). **(G)** Raster plot of generated artificial spike sequences for all 10 trials obtained with the same threshold level of 0.04. **(H)** Peristimulus time histogram (PSTH) for all 10 trials with 0.1ms bin. Time 0 is assigned to the first touch event (start of a collision phase). **(I)** Inter-spike interval histogram for all 10 trials with 0.1ms bin. **(J)** Log-log plot of mutual information rate for the data in G) as a function of a bin size.

This hypothesis of *dispersive vibrational transduction* (DVT) implies that whisker micromotions faster than 500Hz are responsible for *frequency-to-time* transformation of tactile signals and their fast conveyance for subsequent uptake in the neural system.

### Dispersive pre-neuronal processing is biologically plausible

Frequency dispersion used to pre-process sensory inputs prior to encoding into spikes is a common evolutionary strategy for sensory signals uptake that enables repurposing generic neural transduction organs that are otherwise frequency insensitive. For example, biomechanical tonotopy along the cochlea having a graded mass and stiffness along the traveling wave propagation direction provides frequency-to-place transformation of sound (*44*) and induce opening of mechanosensitive membrane channels (*45*) that otherwise are insensitive to frequency. Similarly, glabrous skin vibrations with frequencies up to 800Hz while propagating along the finger skin in primates exhibit strong frequency dispersion of decay rate and propagation speed (*46*), that induce precisely time-aligned spikes in primary afferents (*47*). The DVT hypothesis implies that a whisker is such a dispersive pre-neuronal organ that converts multiple high-frequency eigenmodes excited at the whisker tip during collision, into a time series of vibrational energy bursts arriving at the whisker follicle at different times (Fig.6A). Since whisker is suspended in a collagenous skeleton in a follicle (*22*) that is better matching the whisker impedance, the shockwave is not reflected (as in the case of a whisker glued to aluminum microphone), but rather completely dissipated. However, since this happens only after the shockwave arrives at the follicle, it will not affect the energy dissipation of the modes (and their damping ratios) during their travel along the whisker. It is generally assumed that the mechanosensitive ion channels (*45*) are gated (Fig.6B) by the stimulus force directly and, therefore, can respond with ultrasmall latencies of the order of just a few tens of microseconds (*48*). When the mechanoreceptor membrane potential exceeds a certain threshold then spikes are generated (Fig.6C). Rapidly adapting mechanoreceptors (RA) (i.e. low threshold longitudinal Lanceolate endings (*49*)) are of a particular interest as their spiking is activated by a rapid ramp (whisker angular velocities at least 500°/s (*50*)) of mechanical stimulation and their impulse numbers (spikes per stimulus ramp) are very low, close to just a single spike (*51*) (Fig.6C). Therefore, RA mechanoreceptors are particularly suitable candidates for detecting higher order eigenmode’s wave packets associated with whisker angular velocities up to 5000°/s (Fig.5C). Tracking elastic strain potential energy in the pinned phase, its conversion into kinetic energy during the unpinned phase, and the final transformation into vibrational energy at the collision (Methods) gives an estimate for higher order mode#9 at 3800Hz in Fig.5J delivering about 160nJ to the follicle. It is a hefty 14 orders of magnitude higher than possible sensitivity limit of mechanoreceptors (e.g. 10^-21^J has been reported (*52*)). Primary afferents in rodents TG are also known to produce precisely temporally aligned spikes with exceptionally small jitter of just 10μs (*34*). It is plausible, therefore, that the arrival of individual wave packets of higher order modes to the follicle may be detected by RA.

### High order modes contribute to neural encoding of haptic cues

High frequency rate coding could not be provided by a single mechanoreceptor since neurons in general could not generate spikes at rates faster than 1KHz corresponding to a refractory period of not less than 1ms. However, neurons, and especially RA mechanoreceptors, can generate spikes that are precisely time-aligned to the stimulus presentation with time jitter varied by less than a millisecond from trial to trial(*15, 34, 53*). Instead, following the formulation of the DVT hypothesis, we are proposing here a simple temporal spike encoding model (Fig.6D-H). It is assumed that a sizable population of RA mechanoreceptors is readily available to generate high-frequency population spike train upon arrival of a shockwave bursts even though a single RA receptor will be silent for the duration of its refractory period after firing its very first time-aligned spike. This assumption is well grounded in the neuroanatomy of a vibrissae follicle that has a substantial innervation with a number of mechanoreceptors in the hundreds (e.g., (*54*)). Therefore, our model relies on a population coding as opposed to a single neuron code.

It is also assumed that the RA mechanoreceptors generate a single spike (*51*) when the vibrational energy at the whisker base crosses some threshold power level (both up and down crossings in Fig.6C). For a moderate threshold of 5% the population spike train consists of 20 spikes (Fig.6E) with dense sub-ms inter-spike interval (ISI) within the first 1ms after the collision followed by a sparse spiking for longer times. Each collision event, therefore, generates a unique sequence of timed spikes for the RA population (Fig.6E). Since the power temporal profiles are highly reproducible for repetitive collision events (Fig.6F, see also SI Movies 1, 2, and 5), so are the spike “bar codes” shown in a spike raster plot of Fig.6G for 10 separate whisker sweeps for the same event#2. The resulting peristimulus time histogram (PSTH) in Fig.6H shows distinct well-separated sharp peaks with less than 100μs jitter for each individual peak. While the simple thresholding method produces deterministic spikes, the ISI diagram of Fig.6I shows the presence of some uncertainty mostly determined by trial-to-trial variability of power traces (variability of whisker sweep trajectories resulting in a variability of eigenmode excitation and, therefore, wave packet arrivals).

The number of generated spikes can be tuned by changing a threshold (inset in Fig.6D). Decreasing the thresholding level below 10% results first in a plateau of 20-25 spikes per single collision event (inset in Fig.6D), followed by a fast increase of a spike count above 50 when the threshold is below the background level. Biologically, the threshold tunability can potentially be provided by cortico-thalamic feedback or by adjustment of the impedance mismatch via regulation of a follicle blood flow (*12*).

While the DVT hypothesis considers whisker vibrations, it provides a coding model that is much simpler and with less parameters (Fig.6A-E) than the whisker resonance model (*27, 55*). The latter considers the lowest frequency fundamental mode as a sensory input for spike encoding that can be decoded upstream by the cortico-thalamic phase-locked loops (*56, 57*) for texture discrimination. However, as demonstrated above, the fundamental mode is strongly damped (Fig.3B,D), its frequency is much less affected by the whisker length (Fig.2F,H) than by contact mechanics (*38*), it carries just a fraction of a vibrational energy for both, pole (Fig.S5C, see also (*42*)), and for texture (Fig.4C,G) interactions, and its wavepacket arrives to the follicle the latest with several ms delay.

The DVT hypothesis is consistent with most if not all experimental findings related to verifying the “stick-slip” hypothesis (*18, 28–32*). Various coding schemes have been proposed for interpretation of reverse correlation studies (*14, 15*) of stick-slip induced neural activity including fast variation of whisker curvature (*17, 20*), of angular velocity (*17, 29, 32*), associated angular acceleration peaks (*15, 28*), rotational forces (*15*), or more complex multiplexed products of kinematic and mechanical variables (*58*). The “stick-slip” hypothesis, therefore, implies that single event is not sufficient for rate coding of texture features (and for discrimination), but rather the mean frequency of these events averaged over repetitive swipes. Hence, discrimination of various textures is severely limited since information capacity is reduced by temporal and trial-to-trial averaging.

In contrast, the DVT hypothesis implies that all these variables do not form an independent basis in a multi-dimensional textural feature space, but rather they all are dependent variables of the same process that generates higher-order modes: conversion of elastic strain energy stored in a whisker bending that is released into a kinetic energy of angular whisker acceleration and transformed into high-frequency modal vibrations at collision (Fig.6D). A unique temporal “bar code” (Fig.6E) is generated by each high-speed collision that enables texture discrimination not only with a single whisker in a single sweep, but potentially during just a single collision event.

### Information capacity delivered by higher order modes is in Kb/sec range

To evaluate an upper level of information capacity available at the whisker follicle for the neural uptake after a single collision event, information theoretic methods were employed (*59, 60*) (Methods). The population spike train for each trial was transformed into a binary sequence of 1 (if the spike is present within a given time bin) and 0 (if the spike is absent). Following the Direct Method (DM) (*59*), the Shannon total entropy was determined (*60*) by Eq.8 and noise entropy given the vibrational signal as stimulus was estimated using Eq.9 in Methods. The mutual information (Fig.6J) was then determined by Eq.10 (Methods) as the difference between the total entropy and the noise entropy and is measured in units of bits/sec. For 1ms bin time the mutual information available for spike encoding at the whisker base is 720bits/sec, which is consistent with estimates based on recorded spikes in primary afferents in a somatosensory system (up to 500bits/sec in rodent’s TG (*61*)) and in auditory system (up to 370bits/sec (*62*)). However, when shorter time intervals are considered (time bin size decreased to 200ms), the mutual information is increasing dramatically up to 3000bits/sec (Fig.6J). Further decrease of the bin size down to 50ms sampling limit shows a plateau at around 4000bits/sec. This estimate is indicative of an ultrahigh information capacity that is available for encoding in primary afferents that is at least 10-fold higher than what is usually perceived.

As the spike transduction model above is based on simple thresholding and therefore is deterministic, this estimate does not consider inevitable fluctuations in spike encoding due to stochastic ion channel activation in mechanoreceptors. Hence, the DM estimate provides an upper bound for the mutual information rate. The DM method is, however, based on the assumption that spikes in different bins are not correlated and variance of trial-averaged spiking is assigned to channel noise. However, the temporal “bar code” is not only unique (Fig.6E) for a given collision but is also highly repetitive (i.e., event *a* in Fig.4A-C and in SI Movie.2). Indeed, just a single collision event (e.g., event 2 in Fig.5 and Fig.6) generate modes power profiles with exceptionally low trial-to-trial variability (Fig.6F) for 10 different trials (see also SI Movie.5). The time jitter of all generated spikes for all 10 trials is less than 100μs (Fig.6H) that provides unique information on individual collision event. Texture discrimination then can potentially be done during a single sweep in a single trial without the need for trial-to-trial averaging. In this case the information capacity can be estimated as a number of distinguishable bits in a unique bar-coded word. Assuming the “bar code” length of a few ms (Fig.6F) and 100μs spike jitter (Fig.6H), the information capacity can potentially be boosted up to tens of Kbit/sec.

It has been cautioned that generating spikes with high temporal accuracy is metabolically expensive (*63, 64*), however the DVT hypothesis implies that time coding is needed for just a few ms immediately following the collision event. During this time higher order eigenmodes excited by the most energetic high-speed collision events arrive at the follicle at high group velocity (30-40m/s as in Fig.3H), transduced in RA with short latency into precisely timed spikes (Fig.6E), which are routed to thalamo-cortical circuits via myelinated afferent fibers at matching conduction speeds (10-30m/s (*49, 51*)) ensuring overall ultrasmall latency of sensory uptake. In this scheme the signaling metabolic cost of generating high-bandwidth signals is large while the fixed cost of keeping a signaling system in a state of readiness (that dominates the total cost (*64*)) is low as it is required only for short bursts of activity. Such coding scheme resembles burst-mode optical communications that is considered for large scale datacenters to provide energy-efficient high-bandwidth messaging during short time bursts when circuits are powered up (*65*). Later vibrational events at the whisker follicle defined by arrival of lower order modes can be detected using sparse rate coding that is slower and less metabolically expensive. The overall sensory uptake is multiplexed, therefore, between these slow rate-coded channels and the ultrafast time-coded channels (*50, 57, 58*).

## MATERIALS AND METHODS

### Pre-neuronal processing of haptic sensory cues via dispersive high-frequency vibrational modes

Yu Ding^a^ and Yurii Vlasov * ^a,b,c,d^ All the procedures described in this study were approved by the Institutional Animal Care and Use Committee of University of Illinois Urbana Champaign (protocol #20138), and were performed in accordance with the NIH Office of Laboratory Animal Welfare’s Public Health Service Policy on Humane Care and Use of Laboratory Animals and Guide for the Care and Use of Laboratory Animals. This study is in accordance with the ARRIVE guidelines.

### Whiskers harvesting

We used full grown whiskers from 2 male adult C57BL/6J mice (P56, 30g weight) and a Sprague Dawley male adult rat (P100, 350g weight). Whiskers were plucked from a mystacial pad keeping follicles intact while animals were under general anesthesia by 3% isoflurane inhalation. Shapes of the whiskers were measured under light microscope at high magnification to determine total arc length *S*_*tot*_, diameter of the whisker at the base *R*_*b*_ and at the tip *r*_*t*_, as well as intrinsic curvature (SI Table S1). Intrinsic curvature was defined as a coefficient *A* in the parabolic fitting *A x*^2^of the whisker shape (*20*). Distal radius slope is calculated as *Slope_R_* = (*R_b_ – r_t_)/S_tot_*. For some experiments whiskers were trimmed from the distal side using a sharp razor blade. Resulting total arc length was recorded as well as the change in intrinsic curvature parameter if any. SI Table 1 summarizes measured data from all 4 whiskers.

### Whisker mounting on electret microphone

To measure whisker vibrations during interactions with objects, the electret condenser microphone with built-in op-amp pre-amplifier (Adafruit MAX4466) was used. The covering cloth was removed from the microphone capsule and the follicle end of a whisker was attached directly to the very center of the exposed electret membrane with a nL-scale drop of a glue (Ethyl-2 Cyanoacrylate, Sigma-Aldrich). The glue covered less than 0.02mm^2^ area of the membrane to minimize possible membrane acoustic distortions (See inset in Fig.S1A). Microscopy confirmed that when glued, the proximal base of the whisker was perpendicular to the plane of the membrane. Whisker was attached to the microphone within 24 hours, and all consecutive measurements were done within 48 hours, from the whisker harvesting, respectively. Whisker mechanical properties are believed to be stable for even weeks-long time periods (*21*).

### Scanning system

Microphone was positioned at the center of a rotational scanner driven by a stepper motor (6V Nema-17) through a conveyor belt to minimize motor vibrations (Fig.S1A). The gain of the microphone pre-amplifier was set at a minimal 25X to provide largest dynamic range. The output voltage from the microphone amplifier was digitized at 20kS/s at 16bits (USB-6003, National Instruments). The stepper motor was controlled by a microcontroller (Allego A3967) that was receiving driving voltages at 20kS/s generated with a 12-Bit Analog Output board (PCI-6713, National Instruments). The same driving signals were used to trigger microphone data acquisition. Using the MATLAB Data Acquisition toolbox (R2018b, Mathworks), the analog output and the analog input from the NI devices are digitized and synchronized under the same scanning rate of 20kS/s.

### High-speed videography system

Whisker movements across various objects were simultaneously recorded in two orthogonal planes. The video in the xy plane (Fig.S1B, see also SI Movie.3 and Movie.4) was captured using the overhead 656X600 pixels camera at 1000-1500 fps (Mikrotron EoSens) equipped with a 0.36X telecentric lens that produced 25mmx23mm field-of-view (Edmund Optics, no. 58-257). Video in the yz plane (Fig.S1C) was captured using the side 659X494 pixels camera at 120 fps (Basler acA640) and a 16mm lens that produced 34mmx23mm FOV. Video streams were digitized with a frame grabber (BitFlow Axion) controlled by StreamPix7 multicamera software (NorPix). Each frame was triggered, and the time stamps were generated with less than 1ms jitter. The whole system was illuminated by an overhead LED light (Thor Labs M810L3) powered by DC power supply (Mean Well RS-15-24) and focused with 40 mm focal length aspheric condenser (Thorlabs, ACL5040U).

Slow quasi-steady-state movement of the whole whisker due to scanning at a few Hz frequency can be tracked by extracting individual lossless frames for subsequent manual analysis or using automated tracking with DeepLabCut (*66*). However, the DLC tracking method and, in general, the fast videography, could not be used to characterize high frequency whisker micro-motions beyond 1KHz due to severe limitations in temporal and spatial resolution that results in diminishing dynamic range.

### System calibration

Linear response of the recording system (microphone, amplifier, and ADC converter) is critical for the analysis and the conclusions. Therefore, we performed comprehensive measurements of the linearity for all frequencies of interest starting from 60Hz (frequency of the fundamental vibration mode) up to 10KHz presented in Fig.S2. The dynamic range of the system is evaluated by comparison of the background signal measured with motor movement disabled (Fig.S2A, red) and signal trace with rail-to-rail (0V-3V) span (Fig.S2A, black) measured with a C1 mouse whisker with a total arc length of 22mm swept at 3.5Hz rate against the pole located 2mm apart from the whisker tip. Over one period of sweeping is shown with both concave forward (CF) and concave backward (CB) whisker slip-offs from the pole. The recorded electric power calculated as 20log (*V* − *V*_*off*_) for both signals is shown in Fig.S2B demonstrating over 50dB dynamic range. The offset voltage *V*_*off*_ was measured separately for each trial as a mean voltage at time interval of whisker free vibrations in the air. For example, in Fig.S2A V_off_=1.423V for time interval [0.09s-0.10s]. For all voltage traces in Figs.1-5 the voltage offset was subtracted.

The voltage traces from the microphone were calibrated using the sound and vibration analyzer (977W, Svantek). A speaker (Companion 2 series III, Bose) was used to provide a calibrated input of acoustic field of a given strength at specific frequencies. Sound waves were received by both the sound analyzer and the mounted electret microphone, both located about 50cm away from the speaker. The sound analyzer provides a dynamic analysis of the peak frequency (1/3 octave band) and its field strength in units of dB of Sound Pressure Level (SPL) as 20*l g*_10_(μPa). The magnitude of the harmonic peak in the voltage traces *V* was measured in 1sec recorded clips for each frequency and field strength. Resulting calibration is presented in Fig.S2C. Horizontal axis in this plot is the amplitude of an acoustic harmonic signal accepted by the microphone (in units of sound pressure level, SPL db) and the vertical axis is the electric power of the corresponding recorded voltage signals (in units of 20logV) recorded by the system. These measurements demonstrate that whisker micro-vibrations can be tracked with highly linear response (note the near unity slope of all curves for all frequencies) with a dynamic range spanning 5 orders of magnitude down to the noise floor at -55dB. The system low-signal gain is almost constant across the whole frequency spectrum from 60Hz to 10KHz (note similar x-offset of all the curves).

### Experimental Procedure

During the experiment, the whisker was swiped over an object (pole, grating, sandpaper) located at a fixed distance from microphone (base of the whisker). To mimic the natural whisking motion of rodents in the foveal whisking mode, the scanner was rotated forward and backward at frequencies 3-15Hz to produce 10-15 degrees amplitude sweeps (*6*). Intrinsic curvature of whiskers was aligned with the direction of the xy sweeping plane. The direction of whisking was identified as concave forward (CF) and concave backward (CB) (*20*), consecutively, defined by the intrinsic whisker curvature. Three types of experiments were performed: interaction with a pole, interaction with a textured surface, and interactions with a periodic grating.

#### Interaction with a pole

As observed with video recordings from the overhead and the side view cameras (Fig.S1) the whisker, when moved concave forward (CF), is increasingly bending against the pole (t<0) and is then slipping off the pole at t=0 to vibrate freely in the air. Resulting vibrations produce high voltage signals within the first few milliseconds after the slip-off followed by gradual decay. Complex oscillatory pattern is composed of both low frequency oscillations with a period of a few ms that are dominant at longer times, as well as much faster signals.

Fig.S3A shows voltage traces recorded during interaction of a whisker with a metal wire with 1mm diameter that was placed at varying distances from the whisker base. The characteristic short peak is identified by videography as the last physical contact of the whisker with the pole during slip off. The vibrational energy estimated as an integral under the square of the voltage trace within 30ms time range is increasing exponentially (Fig.S3B) with the pole moving closer to the whisker base. Fig.S3C shows voltage traces recorded after consecutive trimming of the whisker from the intact 21.5mm to 12mm with a constant pole position at 2mm from the whisker tip. Note the multipliers shown for corresponding traces that indicate inverse dependence of the vibrational energy of the whisker length (Fig.S3D). Note that for the 12mm trace the peak voltage exceeds the limits of the op-amp and exhibits rail-to-rail clipping.

#### Interaction with a textured surface

Fig.S4A shows a series of voltage traces recorded with the same whisker while sweeping against sandpapers with different grits roughness (p60 to p400). Signal recorded while the whisker was sweeping in the air is shown for comparison (magenta). Note characteristic peaks at 0 and 180 deg phases that correspond to changing of the sweep direction. Fig.S4B,C shows corresponding distribution and rug plots (Fig.4B) as well as FFT power spectra (Fig.4C).

#### Interaction with a grating

A custom-made aluminum grating with 2mm wide and 1mm deep slots at 4mm period was placed at various distances from the whisker base as shown in Fig.S1.

### Wavelet analysis

The well-known Fast Fourier Transform (FFT) is applicable to the frequency analysis of stationary signals. We are interested, however, in the analysis of fast changes of a signal frequency component. To study the frequency spectrum of whisker micromotions a Complex Morlet Wavelet (CMW) transformation was performed to capture time-varying modes including both low and high frequencies However, due to fundamental uncertainty principle, better time resolution inevitably results in worsening of the frequency resolution.

To achieve balanced time-bandwidth resolution, we used CMW (MATLAB R2018b, Mathworks) with a center frequency of 3Hz (moderate time resolution) and a bandwidth coefficient of 3Hz (moderate frequency resolution). The sampling frequency from the microphone is sampled at 20kS/s, corresponding to Nyquist frequency of 10kHz. When calculating the wavelet coefficients, the frequency domain is partitioned into 5000 steps by an inversely varying scale sequence. The result is a spectrogram of wavelet coefficients with entries corresponding to different frequency components (from 2Hz to 10kHz in log scale) at each sampled time. Wavelet transforms are known to exhibit boundary effects called the cone of influence. To avoid such artifacts, the wavelet transform is performed on a larger time domain (typically 0.1 seconds longer) than the region of interest. On all CWM spectrograms the absolute value of the wavelet coefficients was plotted against time and frequency in a logarithmic scale.

### Fourier filtering analysis

To analyze power evolution of specific modes the CMW method is not optimal as fundamental tradeoff between frequency and time resolution is varying in time and frequency domains. Instead, we used the FFT band-pass filtering applied to voltage traces to analyze how the amplitude changes with time for a single vibrational band (Origin9, Origin Labs). Based on the analysis of a CMW spectrogram (i.e., Fig1B and Fig.1C), the entire 10KHz-wide spectrum was divided into a number of filtering bands centered around the central frequency of corresponding vibrational modes. Lower and upper cutoffs were determined by frequencies of nearby transform amplitude minima that are typically 30dB lower than the amplitude at the peak. Therefore, the bandwidth for each filtering band increases with ascending frequency. As an illustration, the band filters used for analysis of vibrations of a C1 whisker trimmed to 20mm (red curve in Fig.1C) are shown in SI Table S2. Corresponding spectra are shown in Fig.S5A.

### Evaluation of modal power and energy

Since our calibration results of Fig.S2C indicate high linearity of gain throughout the frequency range of interest from 60Hz to 10000Hz, we assume that the voltage induced on the electret membrane of the microphone and then amplified with op-amp is proportional to the power of acoustic waves arriving at the base of the whisker. To estimate the power carried by a particular mode *j*, the offset-corrected voltage traces were bandpass filtered as described above to obtain *V*^*FFT*^(*t*) (i.e., bandpass filtered traces in Fig.S5A). The mode power was evaluated (Fig.S5B) as an envelope of the squared offset-corrected bandpass filtered voltage (*V*^*FFT*^(*t*. The energy of the mode is estimated (Fig.S5C) as an integral under the envelope within a given time interval. Knowing the linear relation between acoustical SPL and the electric power in units of 20log(V) (Fig.S2C), it is possible to convert the received electric power and energy into absolute units of Watts and Joules assuming 100x difference in the microphone active areas (1cm diameter Svantek and 1mm diameter MAX4466) that provides 20dB offset in power density. For example, Fig.S6B right y-axis is expressed in mW, while right y-axis in Fig.S5C in units of mJ. However, this conversion to absolute units can be used cautiously as the calibration does not consider reflection coefficient at the microphone membrane due to acoustic impedance mismatch. Therefore, since our main goal is a comparative study of powers and energies for different modes, whiskers, and excitation conditions, these quantities are presented in arbitrary units unless otherwise specified.

### Evaluation of modal group delay

Modal group delay can be estimated as arrival time of the peak of the power envelope for corresponding FFT bandpass-filtered mode (Fig.S5D). An alternative method to consider spectral signatures of the mode beating due to reflection from the base is shown in Fig.3F,G. Group velocity estimated from group delay is consistent with the results obtained from mode beating as seen in Fig.3H.

### Measurements of damping ratio

Wavelet transformation has varying resolution in time and frequency. For low frequencies, the wavelet is scaled wider for a greater frequency resolution accompanied by worse time resolution. At low frequencies below 200Hz (a typical value of the fundamental frequency), the corresponding scale factor is about 1000 for voltage signals sampled at 20kS/s. The scaled wavelet has a full width at half maximum (FWHM) of 60ms. This hinders the identification of the damping coefficient for the modes below 200Hz using wavelet transform. Above 200Hz the FWHM of the scaled wavelet is below 2ms, capable of resolving the decay. Therefore, to measure damping ratio *ζ* for each vibrational mode we used two separate approaches.

*Frequencies lower than 200Hz.* Identification of the central frequency and the damping coefficient of the fundamental and a few lower order modes below 200Hz from the CMW transformation is hindered by worsened frequency resolution (Δ*j*⁄*j* > 10^−4^). Instead, to identify frequency and damping coefficient of the low order vibrational modes, the FFT bandpass filtered spectrum was fitted with a damped harmonic function (*37*) *V*(*t*) = *A* · *x p*(−2*t*/*τ*) · *n*(*ω*_*n*_(*t* − *t*_*c*_) · *π*/2) as illustrated by a dotted magenta line in Fig.S3. In this particular case the extracted frequency *j* = *ω*_*n*_⁄2*π* is 82±3Hz and damping decay time *τ* is 0.0127 ± 1.9E-4 sec (damping ratio *ζ* is 0.153).

#### Frequencies higher than 200Hz

Damping coefficients of the vibrational modes at frequencies *f* higher than 200Hz (where Δ*j*⁄*j* < 10^−4^) were calculated from the slope of the wavelet spectrogram taken at the central frequency of corresponding mode. It has been shown (*67*) that for the multi-degree-of-freedom (MDOF) systems, the slope of the wavelet coefficient *W*_*g*_*x* for a fixed scale (or dilation) parameter *α*_0_ is a linear function of time *t* and the mode angular frequency *ω*_*n*_:

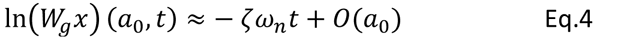

Thus, the damping ratio *ε* = 1⁄*ω*_*n*_*τ* of the mode can be estimated from the slope of the straight line of the wavelet modulus cross-section plotted in a semi-logarithmic scale (decay rate *λ* = 1/*τ*) as shown in Fig3A.

### Analytical and numerical simulations

#### Calculation of natural eigenfrequencies

In assumption of a small damping, the natural undamped frequencies of the modes *f*_*j*_ were obtained from the experimentally measured resonant frequencies *j*^*R*^ by compensating the damping-induced frequency shift following 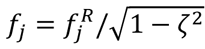. Since measured damping ratios *ζ* are not exceeding 0.35 (see Fig.3B,D) for the fundamental mode and are strongly decreasing for the higher order modes, this correction results in a slight adjustment (below 10%) of the fundamental mode frequency and is negligible for higher order modes.

To fit the natural frequencies of the modes for various truncations of a trimmed whisker (Fig.2) using Euler-Bernoulli model of Eq.1 we assumed that tapering is linear with length (*20*). The variables *β*_*j*_ (*γ*_*j*_) are the dimensionless coefficients for full (truncated) whisker that depend on the eigenmode number and on boundary conditions (*38*). Table S2 shows dimensionless eigenmode coefficients *β*_*j*_ for the first 20 modes calculated for the fixed-free boundary conditions under various whisker truncation using the published MATLAB code (*37*). Truncated ratios are defined as the ratio of the truncated length (*l*) over the total arc length of the whisker (S_tot_). Note, that a fixed boundary condition at the whisker base is assumed here, which is a reasonable assumption for a whisker base glued to the microphone membrane. However, this might be different for a whisker suspended in a follicle (*22*).

### Numerical simulations of dynamical shear forces at the whisker base

We calculated a time-dependent shear force (Fig.2A) generated at the base of the whisker after a slip-off from a pole using the MATLAB code (*37*). The following parameters were used closely matching experimental results for B1 rat whisker trimmed to 29mm: untrimmed total arc length *S*_*tot*_ of 40mm, radius at the base *R*_*b*_=60mm, density 1000kg/m^3^, Young’s modulus 2GPa. For these calculations we assumed a frequency independent damping coefficient α = 430 rad/s for all modes (*37*). Other parameters used in the code: Time_step = 1e-5; Npoints = 10,000; Beta_max = 30,000; Red_factor = 100,000; Freq_limit = 1,000,000; Zpoints = 5000; Xpoints = zpoints.

### Analysis of video recordings

#### Synching video frames

Since the scanner was actuated through a conveyor belt to minimize vibrations, the lag in back and force movements resulted in a 5ms jitter. To better synchronize video frames to the microphone signal, an additional procedure was used. Within each sweeping period, the video frame that captures the whisker first touch (or collision) event was manually identified and then time-aligned to the onset of a large voltage peak in the microphone signal recording. Such peaks (i.e., Fig.S3 at t=0) form a periodic pattern in a microphone signal recorded during consecutive sweeps and are identifiable with better than 0.25ms precision. An example is shown in Fig.S6C with frames time-synchronized to microphone recording of Fig.S6D.

#### Post-processing of video recordings

Individual frames from recorded video were post-processed as illustrated in Fig.S6. A single background image that was shot when whisker was outside the camera field of view, was subtracted from each original frame in a video (Fig.S6A) resulting in a high contrast processed frame (Fig.S6B). An example of a processed video is in SI Movie.3 and Movie.4.

#### Evaluation of forces at the whisker base

The force at the base of whisker was extracted from the whisker curvature following established methods (*9*). In each post-processed frame the coordinates along the whisker were segmented by thresholding and averaging over each y-coordinate. Then a 5^th^ order polynomial was fitted to both x and y coordinates for arc length parametrization. The whisker curvature κ_p_ was estimated by fitting a second order polynomial to a whisker segment centered at one-third of a total whisker arc length. For the example shown in Fig.S6B, a 2mm whisker segment centered at 19mm arc length was used. The intrinsic whisker curvature was subtracted.

#### Tip Acceleration

From each post-processed frame, the contact point between the tip of the whisker and the object was identified. The contact angle was defined as the clockwise angle between the horizontal axis and the contact point. The x-y spatial displacement of a 0.5mm whisker segment next to a contact point taken at a constant arc length was tracked through all frames. The spatial displacement was converted to angle velocity and angle acceleration using calculated contact angles and frames timestamps.

### Estimation of elastic strain energy

Assuming that shear and torsional strain potential energies are negligible, the elastic strain potential energy *U* accumulated during the pinned phase can be estimated (*68*) as,

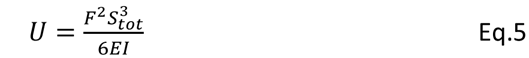

where *F* is the force applied to a whisker. The latter can be estimated from the maximum curvature *κ*_*p*_ at the beginning of the unpinned phase (*9*) at a point *p* along the whisker arc,

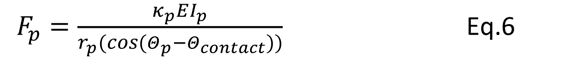

where *I*_*p*_ = *π r* *α*^4^⁄4 the second moment of inertia at point *p*, *α* is a whisker radius at *p*, *r*_p_ is the arm of the force at p, *θ*_*p*_ is an angle of vector connecting *p* to the site of whisker-object contact, and *θ*_*p*_*n t α*_*contact*_ is an angle of a whisker at contact.

Combining Eqs.S2 and S3 we arrive to the following equation,

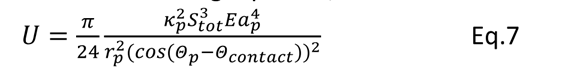

### Estimation of vibrational energy delivered by a high-frequency mode

To estimate how much energy is delivered to the follicle by high-frequency modes, it is instructive to check for the energy conservation in the three-phase pin-unpin-collision process. For the event 2 in Fig.5, the elastic strain potential energy *U* accumulated during the pinned phase can be estimated from the maximum curvature at the beginning of the unpinned phase (Eq.7), assuming that shear and torsional strain potential energies are negligible. Part of this energy, equal to the drop of a curvature *Δ κ*_*p*_=0.26mm^-1^ at the transition to the unpinned phase in Fig.5, is transformed into a kinetic energy that can be estimated from Eq.7 as 6mJ. The upper limit of the impact energy released at the collision is equal to rotational kinetic energy *K* = *I*_*p*_*ω*^2^/2. Assuming that rotational velocity at collision *ω*=50rad/s, this gives the impact energy of about 4mJ. This corresponds to over 65% of the kinetic energy in the unpinned state showing reasonable conversion efficiency. The rest is most likely dissipated in a series of subsequent periodic collisions with diminishing amplitudes (see oscillations in acceleration in Fig.5C and Fig.5J also visible in the SI movie4). Therefore, the energy of a higher order mode#9 at 3800Hz that carries about 10% (Fig.6J) of a total released impact energy, is 400nJ. While arriving at the base, the energy is attenuated by 4dB (from damping in Fig.3A) down to 160nJ. An alternative estimate based on integrating the calibrated squared voltage trace of Fig.6D provides the total vibrational energy at the base as 2mJ and assigns similar 200nJ (10% energy fraction) to the mode#9. While for a whisker glued to a microphone most of this energy is reflected back due to large acoustical impedance mismatch, for a whisker in-vivo most of this energy is absorbed within a follicle due to much closer matching of impedances.

### Estimation of mutual information carried by high-frequency vibrations

Artificial spike sequence of Fig.6E was generated by setting a threshold for the squared offset-corrected voltage (Fig.6D) to 0.04. Spikes were generated whenever the signal crossed a given threshold. To avoid unwanted digital noise during thresholding procedure the voltage traces were up-sampled from original 20Ks/s to 100Ks/s by linear extrapolation between neighboring datapoints. The artificial spike sequence (Fig.6G) were generated for 10 nominally identical sweeping trials in Fig.6H (see also SI Movie.5) using the same threshold. The inter-spike interval (ISI) of Fig.6I was obtained for each trial by counting the temporal lags between all possible pairs of time bins with spikes and then averaged between all trials.

The spike sequence was digitized to 1 bit words by assigning 1 if spikes are present within a time bin Δt and 0 if absent (*60*). The conditional probability of observing a spike at each time bin *p*(1|_*t*_) was calculated by averaging the digitized artificial spike sequences over all 10 trials during a time window of *T*=5ms. Probability of not observing a spike at the same time is *p*(0|_*t*_) = 1 − *p*(1|_*t*_). The probability *p*(*i*) was determined by averaging

*p*(*i*|_*t*_) over all time bins. The total Shannon entropy S_total_ associated with bin size Δt could be determined as (*60*) :

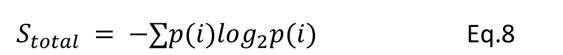

where the sum is over all possible values of words (0 and 1 in this case).

The noise entropy *S*_*noice*_ associated with the time-dependent stimuli is estimated via the Direct Method (*59, 60*) assuming that stimuli in each time bin are independent. The noise entropy of the spike sequence given the vibrational signal as stimulus was estimated by averaging entropy over all time bins:

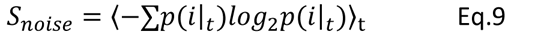

The average mutual information for a time bin Δt was then determined by the difference in entropies (*60*):

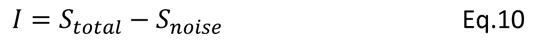

The information rates per time bin Δt and per single spike were calculated by dividing the average mutual information *I* by the time bin Δt or by mean firing rate, correspondingly.

Since information rate estimated by DM is known to be biased due to noise entropy being sampled from a limited number of trials (*69*), the DM results were checked with a jackknifed method (JK) (*70*) that estimates entropy of all but the j-th trial out of N trials. The results of DM and JK mutual information rates for 5 and 10 trials differ by less than 3% presumably due to very low trial-to-trial variability. This is consistent with estimated sampling bias of 1% by quadratic expansion (*59*) looking at the dependence of entropy on the fraction of trials (from 10 trials down to 5 trials).

## SUPPLEMENTARY INFORMATION

**Fig.S1.**
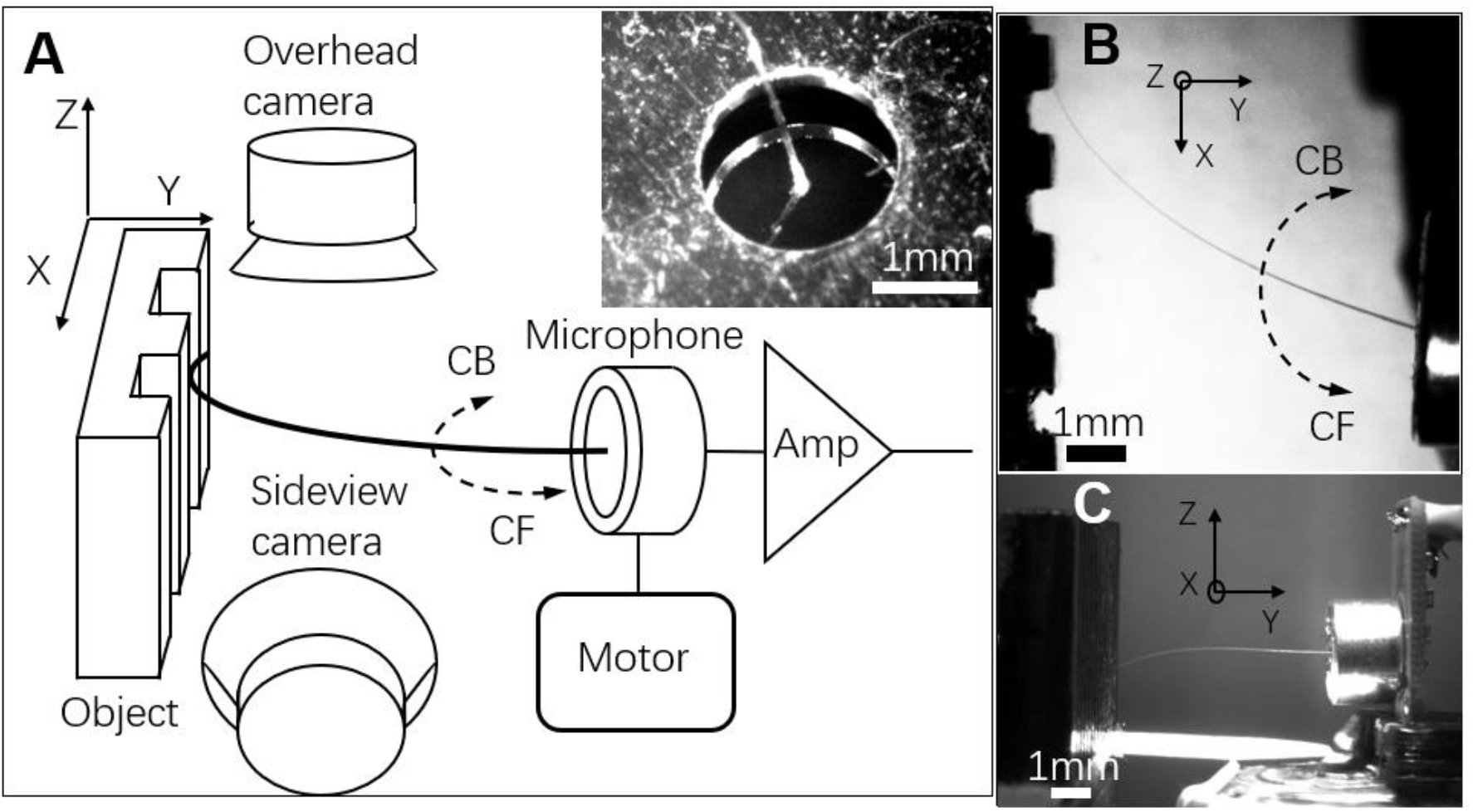
Whisker micromotions acquisition system. **(A)** Schematic of the experimental setup. Inset: Photograph of a whisker glued to the electret membrane of electret condenser microphone. Vibrations induced on the membrane was transformed into voltage traces, amplified, and recorded. The microphone was swept in the xy plane using a stepper motor. Whisker intrinsic curvature was oriented to align with the sweeping plane thus defining convex forward (CF) and convex backward (CB) directions. An object (here a grating) was placed at y distance from a whisker base, deflecting the whisker during sweeping motion. **(B)** A frame image of a whisker micromotions in the *xy* plane captured with the overhead camera. **(C)** A frame image in the *yx* plane captured with the side view camera.

**Fig.S2.**
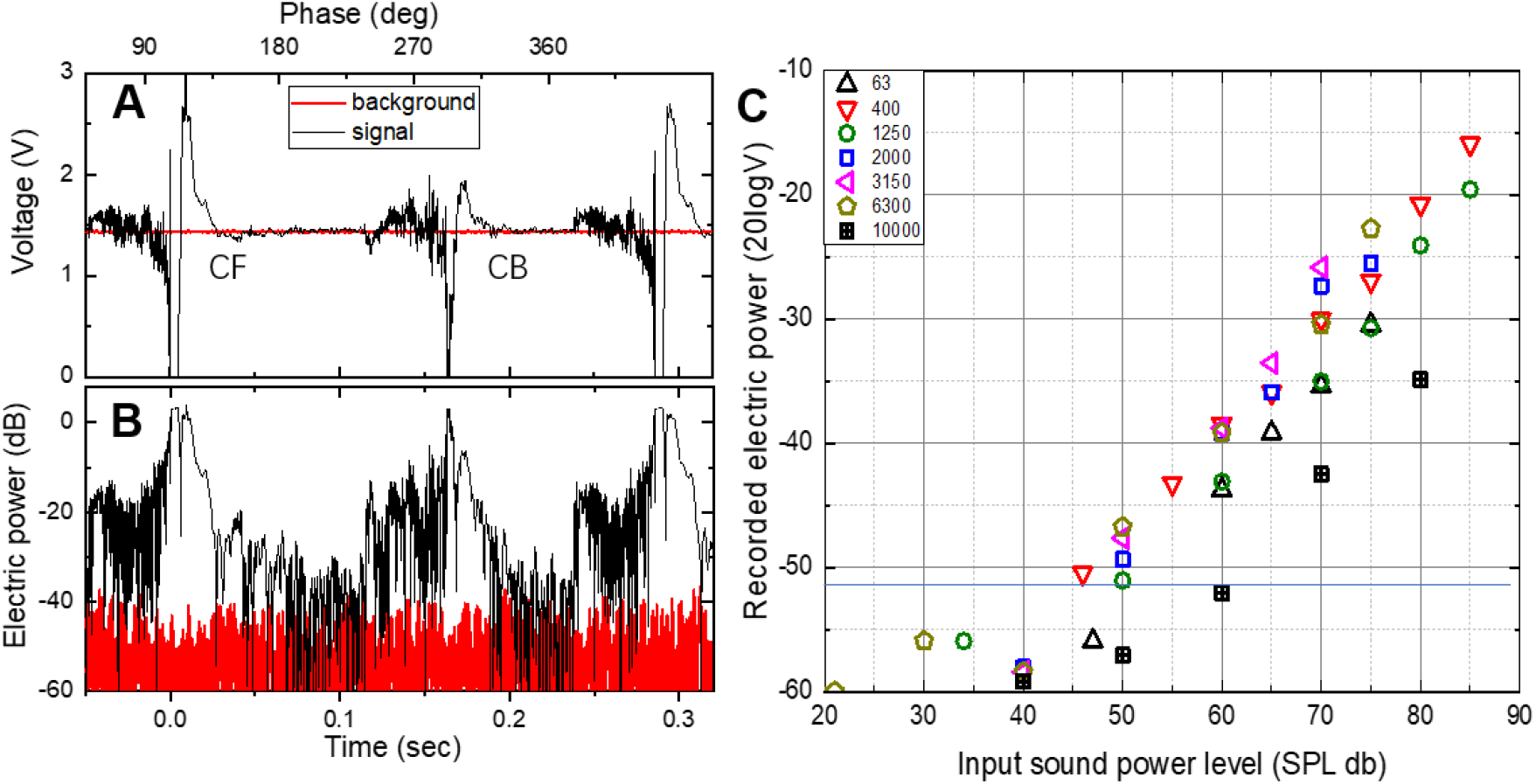
Calibration of acoustic system. **(A)** Voltage trace recorded with a C1 mouse whisker (black) with a total arc length of 22mm swept at 3.5Hz rate against the pole located 2mm apart from the whisker tip. For comparison, a background trace (red) without motor movement is recorded. **(B)** Electric power calculated from voltage traces for signal (black) and background (red) demonstrating over 50dB dynamic range. **(C)** The electric power (dB) from the electret microphone with attached whisker is plotted as a function of the acoustic field strength (SPL dB). Different symbols with different colors represent calibration at different frequencies (63-10,000Hz).

**Fig.S3.**
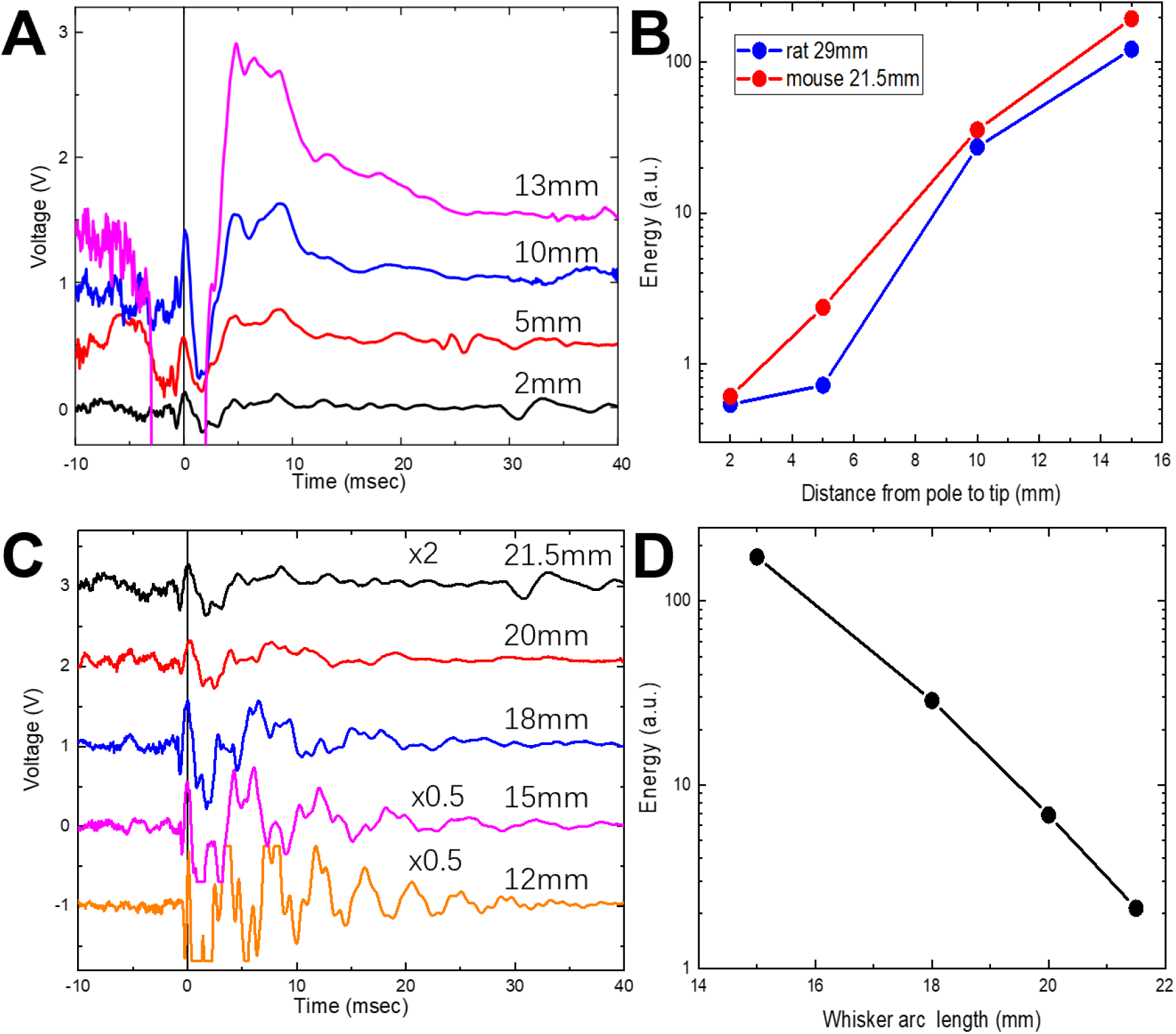
Interaction with a pole. **(A)** Voltage traces recorded with a C1 mouse whisker with a total arc length of 21.5mm swept at 3.5Hz rate against the pole located at different distances from the whisker tip. Traces are shifted vertically by 0.5V with respect to each other for clarity. **(B)** Energy of vibrations calculated as an integral under the squared voltage traces in A) for [0..30] ms time interval. **(C)** Voltage traces recorded with a C1 mouse whisker that is consecutively trimmed form initial 21.5mm to 12mm total arc length. The distance between a pole and the whisker tip is kept constant as 2mm. Traces are shifter vertically by 1V with respect to each other for clarity. Voltage traces for 12, 15, and 21.5mm are scaled with the multipliers shown. Note amplitude clipping of 12mm voltage trace due to saturation of an op-amp amplifier. **(D)** Energy of vibrations calculated as an integral under the squared voltage traces in C) for [0..30] ms time interval.

**Fig.S4.**
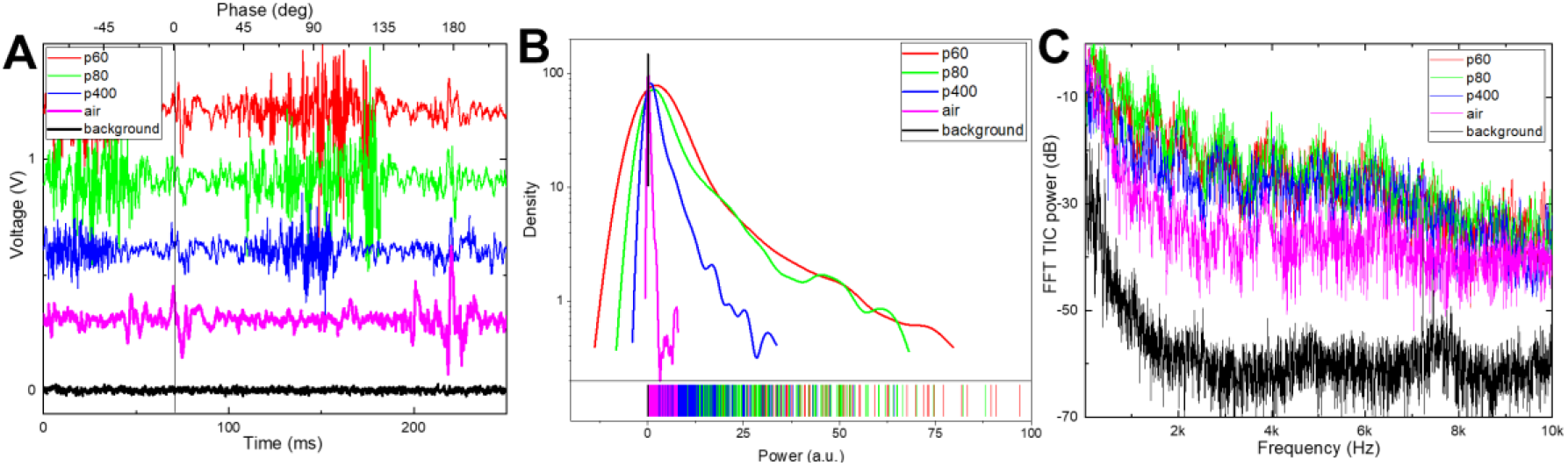
Interaction with a surface. **(A)** Voltage traces recorded with a C1 mouse whisker trimmed to a total arc length of 21.5mm swept at 3.5Hz rate against the sandpapers with different grit numbers. The sandpapers are located at 2mm from the whisker tip. Traces are shifted vertically by 0.25V with respect to each other for clarity. Traces for scanning of a whisker in the air (magenta) and the background noise level with scanning stopped (black) are shown for comparison. **(B)** Distribution and rug plots for traces in A). **(C)** FFT power spectra for voltage traces in A).

**Fig.S5.**
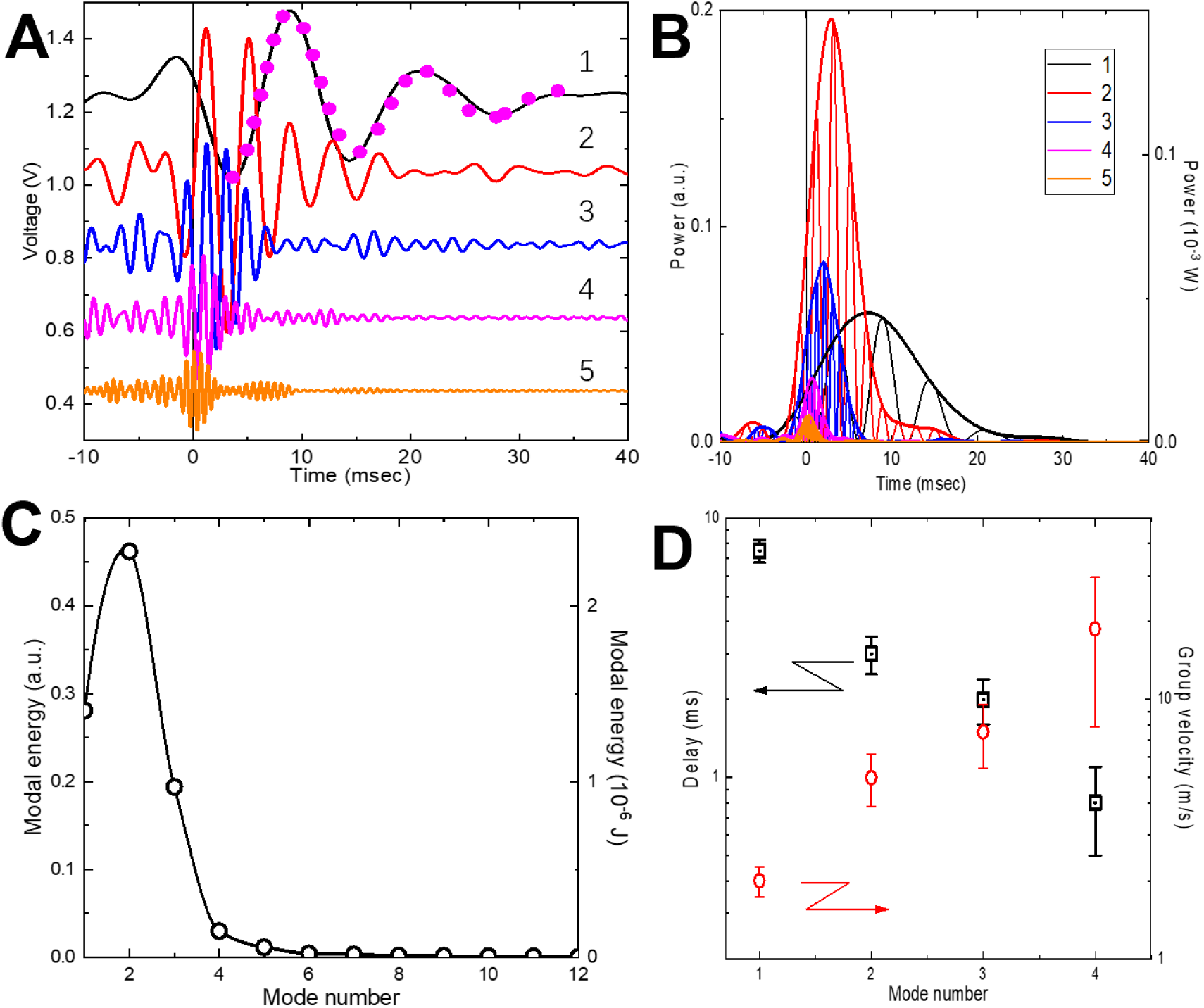
FFT bandpass filtering procedure. **(A)** Voltage traces recorded with a C1 mouse whisker (black) with a total arc length of 20mm swept at 3.5Hz rate against the pole located 2mm apart from the whisker tip. Traces marked 1 through 5 correspond to FFT-filtered bands according to filters cutoffs in the Table S1. Filtered spectra are shifted vertically by 0.2V for clarity. Dotted magenta curve on top of the spectrum 2 is a fitting using a damped sinusoidal function. **(B)** Plot of squared voltages in A) and their envelopes. Modal power is proportional to the magnitude of the envelope. **(C)** Modal energy calculated as an integral under the envelope over a [-5..30] ms interval. Note that second order mode has the highest power. **(D)** Modal group delay and group velocity calculated at the maxima of the power envelopes for corresponding modes in B).

**Supplementary Table S1.** Measured parameters of whiskers.

| # | Whisker | R<sub>b</sub><br>($\mu\text{m}$) | S<sub>tot</sub><br>(mm) | Slope<sub>R</sub><br>($\times 10^{-3}$) | Bending<br>(A) |
| --- | --- | --- | --- | --- | --- |
| 1 | Rat B1 | 60 | 40.0 | 1.45 | 1.0 |
| 2 | Mouse C1 | 37 | 21.4 | 1.53 | 1.9 |
| 3 | Mouse C2 | 35 | 24.1 | 1.48 | 1.1 |

**Supplementary Table S2.** Example parameters for the FFT bandpass filtering.

| FFT band<br>number | Mode central<br>frequency (Hz) | Lower cutoff<br>(Hz) | Upper cutoff<br>(Hz) |
| --- | --- | --- | --- |
| 1 | 90 | 10 | 150 |
| 2 | 270 | 150 | 400 |
| 3 | 550 | 400 | 700 |
| 4 | 1000 | 700 | 1200 |
| 5 | 1600 | 1200 | 1800 |
| 6 | 2300 | 1900 | 2600 |
| 7 | 3300 | 2600 | 3700 |

**Supplementary Table S3:** Dimensionless eigenmode coefficients for truncated conical beam.

|  | 0 | 0.1 | 0.2 | 0.3 | 0.4 | 0.5 | 0.6 | 0.7 | 0.8 | 0.9 |
| --- | --- | --- | --- | --- | --- | --- | --- | --- | --- | --- |
| 1 | 8.72 | 8.89 | 9.68 | 11.24 | 13.91 | 18.50 | 26.99 | 45.19 | 96.38 | 367.37 |
| 2 | 21.15 | 23.06 | 28.73 | 38.04 | 52.96 | 78.19 | 125.31 | 228.39 | 526.42 | 2155.03 |
| 3 | 38.45 | 45.83 | 62.24 | 87.37 | 127.05 | 194.32 | 320.84 | 600.17 | 1415.76 | 5918.86 |
| 4 | 60.68 | 78.40 | 111.32 | 160.25 | 237.07 | 367.25 | 612.53 | 1155.28 | 2744.07 | 11539.78 |
| 5 | 87.83 | 121.19 | 176.29 | 257.04 | 383.43 | 597.56 | 1001.23 | 1895.30 | 4515.27 | 19035.68 |
| 6 | 119.92 | 174.36 | 257.29 | 377.85 | 566.24 | 885.31 | 1486.99 | 2820.22 | 6729.15 | 28405.39 |
| 7 | 156.94 | 237.98 | 354.37 | 522.74 | 785.52 | 1230.55 | 2069.85 | 3930.07 | 9385.75 | 39648.10 |
| 8 | 198.90 | 312.08 | 467.55 | 691.71 | 1041.31 | 1633.28 | 2749.81 | 5224.86 | 12485.10 | 52766.52 |
| 9 | 245.80 | 396.70 | 596.85 | 884.78 | 1333.61 | 2093.53 | 3526.88 | 6704.60 | 16027.19 | 67757.95 |
| 10 | 297.64 | 491.83 | 742.27 | 1101.96 | 1662.43 | 2611.28 | 4401.07 | 8369.29 | 20012.03 | 84623.30 |
| 11 | 354.43 | 597.49 | 903.83 | 1343.25 | 2027.76 | 3186.56 | 5372.39 | 10218.95 | 24439.62 | 103362.57 |
| 12 | 416.18 | 713.69 | 1081.53 | 1608.66 | 2429.63 | 3819.35 | 6440.83 | 12253.56 | 29309.97 |  |
| 13 | 482.90 | 840.43 | 1275.37 | 1898.19 | 2868.01 | 4509.66 | 7606.40 | 14473.13 | 34623.07 |  |
| 14 | 554.59 | 977.71 | 1485.35 | 2211.83 | 3342.92 | 5257.49 | 8869.09 | 16877.66 | 40378.92 |  |
| 15 | 631.27 | 1125.54 | 1711.47 | 2549.60 | 3854.36 | 6062.85 | 10228.91 | 19467.16 | 46577.53 |  |
| 16 | 712.96 | 1283.92 | 1953.74 | 2911.49 | 4402.33 | 6925.73 | 11685.86 | 22241.61 | 53218.90 |  |
| 17 | 799.70 | 1452.84 | 2212.16 | 3297.50 | 4986.82 | 7846.13 | 13239.93 | 25201.03 | 60303.02 |  |
| 18 | 891.40 | 1632.31 | 2486.72 | 3707.64 | 5607.85 | 8824.05 | 14891.14 | 28345.40 | 67829.90 |  |
| 19 | 988.19 | 1822.33 | 2777.43 | 4141.89 | 6265.40 | 9859.50 | 16639.47 | 31674.74 | 75799.53 |  |
| 20 | 1090.05 | 2022.91 | 3084.29 | 4600.27 | 6959.48 | 10952.47 | 18484.93 | 35189.05 | 84211.93 |  |

## Supplementary Movies

**SI Movie 1: Spectral analysis of whisker interaction with a pole.**

Voltage trace (top) of a C1 21.5mm mouse whisker swept over a pole at 10mm distance from the tip. Video (bottom) is composed of 10 CMW spectrograms calculated from 10 consecutive sweeps. Note highly reproducible regular pattern of modal vibrations at times corresponding to a whisker slipping off the pole (downward red arrows) and collision with the pole (upward arrows). Note also, that at the first touch the eigenmodes pattern corresponds to a shorter whisker arc than for a slip-off. Corresponding upward chirp in eigenmodes frequencies is schematically shown by white dashed lines.

**SI Movie 2: Spectral analysis of whisker interaction with a textured surface.**

Voltage trace (top) of a B1 40mm rat whisker swept over a sandpaper with p60 grit value located at 2mm distance from the tip. Video (bottom) is composed of 10 CMW spectrograms calculated from 10 consecutive CF sweeps. Note highly reproducible regular pattern of modal vibrations at times corresponding to a whisker slipping off the sandpaper surface (downward red arrow) and first contact with the sandpaper (upward arrow). Events *a*, and *b* discussed in relation to Fig.4A,B are marked by downward red arrows.

**SI Movie 3: Whisker swipe over grating – Fig.S6.**

Background-subtracted 1000fps video of a mouse whisker scanned over a 2mm pitch grating (same results as in Fig.S6). Note blurring of the images when whisker is vibrating at frequencies beyond 1000Hz.

**SI Movie 4: High-speed collisions during sweeping over a grating – Fig.5**.

Background-subtracted video of a C2 mouse 24.1mm long whisker during a single CB sweep over a grating recorded at 1460fps (same results as in Fig.5). Image of the whisker (thinner bottom trace) is accompanied by an image of the whisker shadow projected on a screen (thick upper trace). Note, that the view of a whisker tip is blocked by a grating teeth. The position of a teeth at the plane of a sweep (left) and the microphone base (right) are shown by rectangular shapes. Events marked 1, 2, and 3 correspond to interaction with individual grating teeth as in Fig.5. Overlaid are the whisker shapes extracted from consecutive 160 frames.

**SI Movie 5: Spectral analysis of interaction of a whisker with a grating – Fig.5**.

Typical voltage trace (top) of a mouse C2 whisker with 24.1mm arc during a single CB sweep over a grating. Events marked 1, 2, and 3 correspond to interaction with individual grating teeth as in Fig.5. Video (bottom) is composed of 10 CMW spectrograms calculated from 10 consecutive CB sweeps. Time for each spectrogram is aligned to the start of event 2.

